# Environmental filtering drives cryptic diversity and shifting interaction networks across lake, ephemeral pond, and desiccated microbial mat communities in the Untersee Oasis, East Antarctica

**DOI:** 10.64898/2026.02.16.706179

**Authors:** L. Vimercati, I. Chakrabarti, A. Lindley, C. Greco, D. Andersen, A.D. Jungblut

## Abstract

Antarctic lakes and surrounding habitats are natural laboratories for studying microbial ecology in extreme environments, yet assembly and dispersal across these systems remain poorly understood. We analyzed bacterial (16S rRNA) and eukaryotic (18S rRNA) communities across benthic lake mats, pond mats, and desiccated mats around Lake Untersee, East Antarctica. Both domains showed significant environmental differentiation with stochastic assembly dominating, but eukaryotes exhibited stronger habitat-specific structuring. Core microbiomes revealed contrasting strategies: bacteria showed broad colonization with few dominant taxa, while eukaryotes maintained smaller cores anchored by *Adineta vaga*. Cyanobacterial indicators demonstrated complete strain-level turnover despite uniform phylum dominance. Dispersal patterns diverged strongly: prokaryotes exhibited substantial cross-habitat exchange (28-61%) while eukaryotes showed minimal dispersal (2-16%). Distance-decay relationships revealed significant spatial structuring (bacterial r=0.316-0.70; eukaryotic r=0.43-0.84). Cross-domain networks shifted from facilitative in ponds (65% positive) to competitive under desiccation (54% negative), suggesting resource limitation overrides facilitation at extremes. Functional predictions highlighted habitat specialization: lakes enriched in biogeochemical cycling, desiccated mats in stress tolerance, ponds in CRISPR genes. These findings reveal divergent strategies shaped by environmental filtering, dispersal limitation, and viral interactions, with implications for ecosystem connectivity and resilience.

**Importance:** Antarctic lake ecosystems serve as natural laboratories for understanding microbial community assembly under extreme conditions, yet connectivity between aquatic and terrestrial habitats remains poorly characterized. This study reveals fundamentally divergent ecological strategies between bacterial and eukaryotic communities across Lake Untersee’s contrasting environments. While bacteria maintain broad dispersal networks (28-61% cross-habitat exchange) with strain-level habitat specialization, eukaryotes exhibit strong dispersal limitation (2-16% exchange) despite occupying adjacent habitats. Most strikingly, cross-domain interaction networks shift from predominantly facilitative in ephemeral ponds (65% positive associations) to competitive under desiccation stress (54% negative associations), directly challenging the stress gradient hypothesis that predicts cooperation dominates in extreme environments. Combined with habitat-specific functional specialization—from biogeochemical cycling in lakes to stress tolerance in desiccated mats—these findings provide a mechanistic framework for predicting ecosystem connectivity, resilience, and responses to climate-driven habitat transitions in polar regions.

## Introduction

Perennially ice-covered lakes across regions of continental Antarctica represent some of Earth’s most stable, yet extreme, aquatic ecosystems. Among these, Lake Untersee in Queen Maud Land stands out as a globally unique water body due to its exceptional hydrochemistry and the complex microbial communities it hosts (Andersen et al. 2011). The lake is characterized by poorly buffered, low-DIC waters with high pH (∼10.4) in its upper water column, which is consistent with its long-term hydrological steady state and extremely low TIC/TOC concentrations (Marsh et al. 2020), supersaturation with oxygen down to approximately 70 meters, and a substantial, permanently stratified anoxic sub-basin containing extremely high methane concentrations. Bordered and fed by the Anuchin Glacier at its northern edge, Lake Untersee provides an ideal model for studying cryo-lacustrine interactions (Weisleitner et al. 2019), with its tightly sealed hydrological regime defined by the lake’s energy and water balance (Faucher et al. 2019). While the benthic and water column communities of Lake Untersee have been relatively well-characterized (Koo et al. 2017; Greco et al. 2020, 2024), understanding of connectivity between the lake and surrounding terrestrial habitats remains limited. The terrestrial landscape harbors diverse microbial niches, including seasonal ponds and desiccated microbial mats, which have received little scientific attention despite their potential ecological significance. Recent mapping and geochemical characterization of these ponds (Faucher et al. 2021a) revealed strong spatial heterogeneity and sensitivity to climate forcing, suggesting that they may serve as important reservoirs of microbial diversity and function as sources or sinks for dispersal to and from the lake ecosystem. Antarctic meltwater ponds demonstrate substantial regional connectivity, with local geochemical conditions driving community structure while maintaining dispersal networks across considerable distances (Archer et al. 2015). Such habitat connectivity becomes increasingly important for understanding Antarctic ecosystem responses to climate change. Recent evidence suggests that external inputs, including cryoconite communities from the Anuchin Glacier, may contribute substantially to lake benthic mats (Weisleitner et al. 2019), yet the major biotic sources for the lake ecosystem remain largely unknown. Additionally, Lake Untersee has been established as a planetary analog for early Mars and icy worlds such Enceladus (McKay et al. 2017), where CO₂-limited, nearly carbonate-free lakes can nonetheless maintain productive benthic communities and undergo episodic carbon enrichment, exemplified by the recent glacial lake outburst flood from Obersee (Faucher et al. 2021b). To address the knowledge gap regarding connectivity between Lake Untersee and its surrounding terrestrial habitats, we conducted a comprehensive multi-domain analysis of microbial communities across three distinct environments: benthic lake mats, surrounding pond mats, and desiccated mats that emerged through the lake ice surface. Using both prokaryotic (16S rRNA gene) and eukaryotic (18S rRNA gene) community profiling, we examined community composition, indicator species, core microbiomes, assembly processes, source-sink relationships, co-occurrence networks, and functional potential to test several key hypotheses. We hypothesized that:

1. Environmental filtering would drive distinct bacterial and eukaryotic community composition across lake, pond, and desiccated habitats, with stronger differentiation in eukaryotes due to greater environmental sensitivity.
2. Stochastic assembly processes would dominate both domains across all environments, but eukaryotic communities would show stronger environment-specific assembly patterns than bacterial communities.
3. Cross-domain association networks would shift from sparse (lake) to facilitative (pond) to competitive (desiccated) following the stress gradient hypothesis predictions (Bertness and Callaway 1994).
4. Source-sink connectivity would be higher for bacteria than eukaryotes, with dispersal occurring primarily among surface habitats (desiccated and pond mats), and desiccated mats expected to be the dominant source due to greater exposure and wind-driven export.
5. Functional specialization would emerge across environments, with lake communities enriched in biogeochemical cycling functions while desiccated communities would show enhanced stress tolerance capabilities.

## Material and Methods

### Study sites and sampling

Benthic microbial mats were collected from Lake Untersee using detailed methods described in Greco et al. 2020. Briefly, scientific divers utilizing SCUBA collected samples from the southern basin at 20 m depth in November 2011 (austral summer) using sterile blades and sterile 50 ml conical tubes. Three distinct macroscopic mat morphologies were sampled: flat, cone, and pinnacle shaped microbial mats, along with a loose mesh of darkly pigmented cyanobacteria filaments from mat surfaces. Flat mat samples were collected from areas between cones and pinnacles. In the field laboratory, pigmented upper laminae (less than 1 mm thickness) were separated from lower non-pigmented sediment-rich portions using sterile blades and rinsed in sterile deionized water prior to DNA extraction. Pond mats were collected in December 2019 from multiple ponds in the vicinity of Lake Untersee. Desiccated mats originated from benthic lake mats that became buoyant and surfaced through the ice due to gas bubble accumulation from microbial metabolism. These detached mat fragments subsequently underwent natural freeze-drying as they traveled across the ice surface, experiencing prolonged exposure to cold, arid Antarctic conditions that removed moisture content while preserving microbial community structure. Samples were stored at −20°C after collection and maintained at −80°C at the Natural History Museum of London until processing. Representative samples of benthic, pond, and desiccated mats used in this study are shown in Fig. 1A. Sampling coordinates for both pond and desiccated mat collections are listed in Table 1 and locations are shown in Fig. 1B. The present study combines these newly collected pond and desiccated mats with the previously published benthic mat dataset from Greco *et al*. 2020 (BioProject PRJNA638357) to enable a unified comparative analysis.

**Figure 1.**
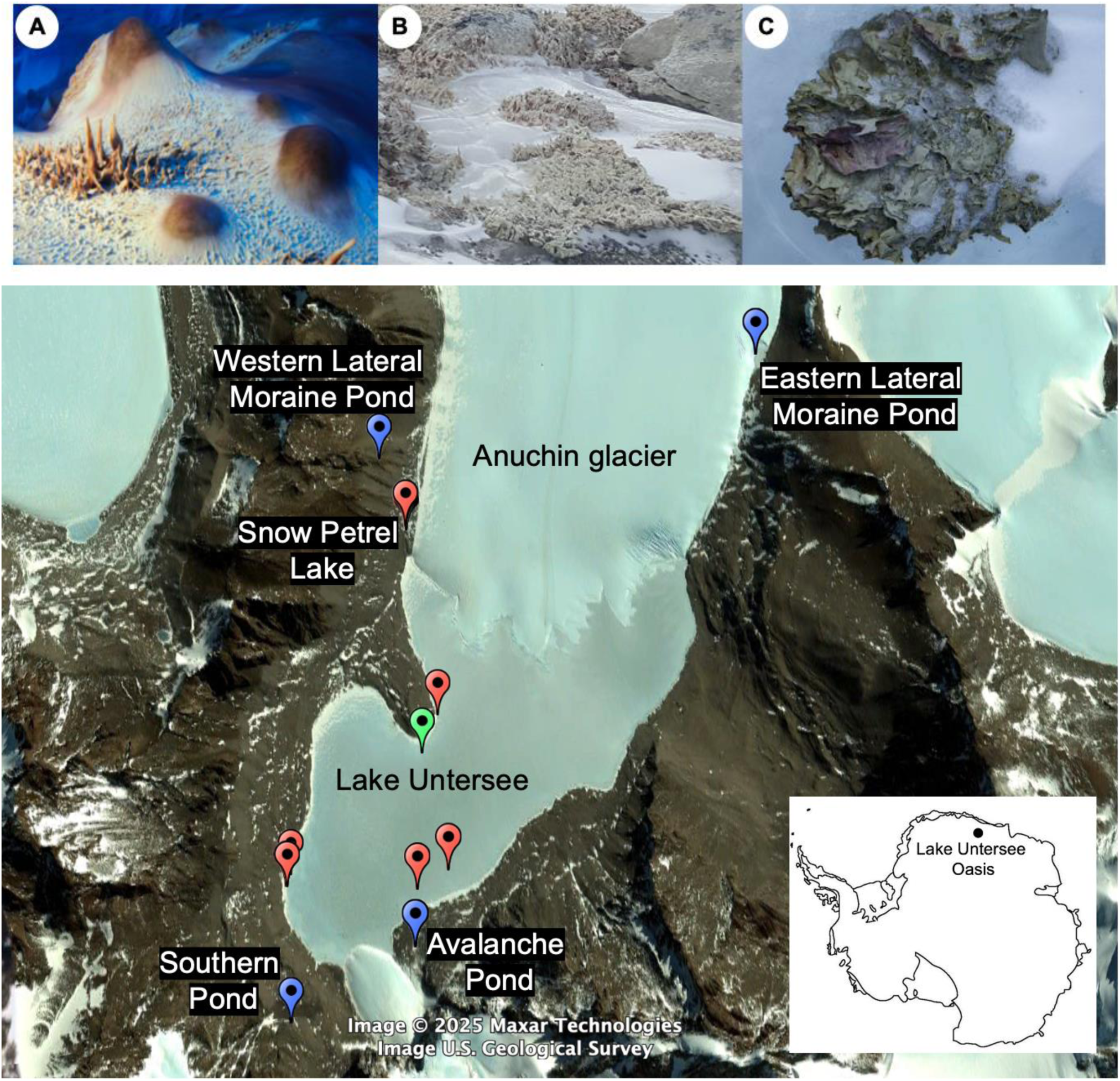
Top: Microbial mat habitats at Lake Untersee, Antarctica. (A) Lake benthic mats at 20 m depth showing flat, cone, and pinnacle morphologies characterized by darkly pigmented cyanobacterial filaments. (B) Pond mats from shallow ponds near Lake Untersee. (C) Desiccated mats derived from buoyant lake mats that surfaced through the ice and underwent natural freeze-drying during transport across the ice surface. **Bottom: Sampling locations at Lake Untersee, Antarctica.** Satellite imagery showing the distribution of sample collection sites across the study area. Green pin indicates the SCUBA dive site where lake benthic mats were collected at 20 m depth. Blue pins mark pond sampling locations. Red pins indicate collection sites of desiccated mats on the ice surface. Satellite imagery: © 2025 Maxar Technologies, U.S. Geological Survey.

**Table 1.**
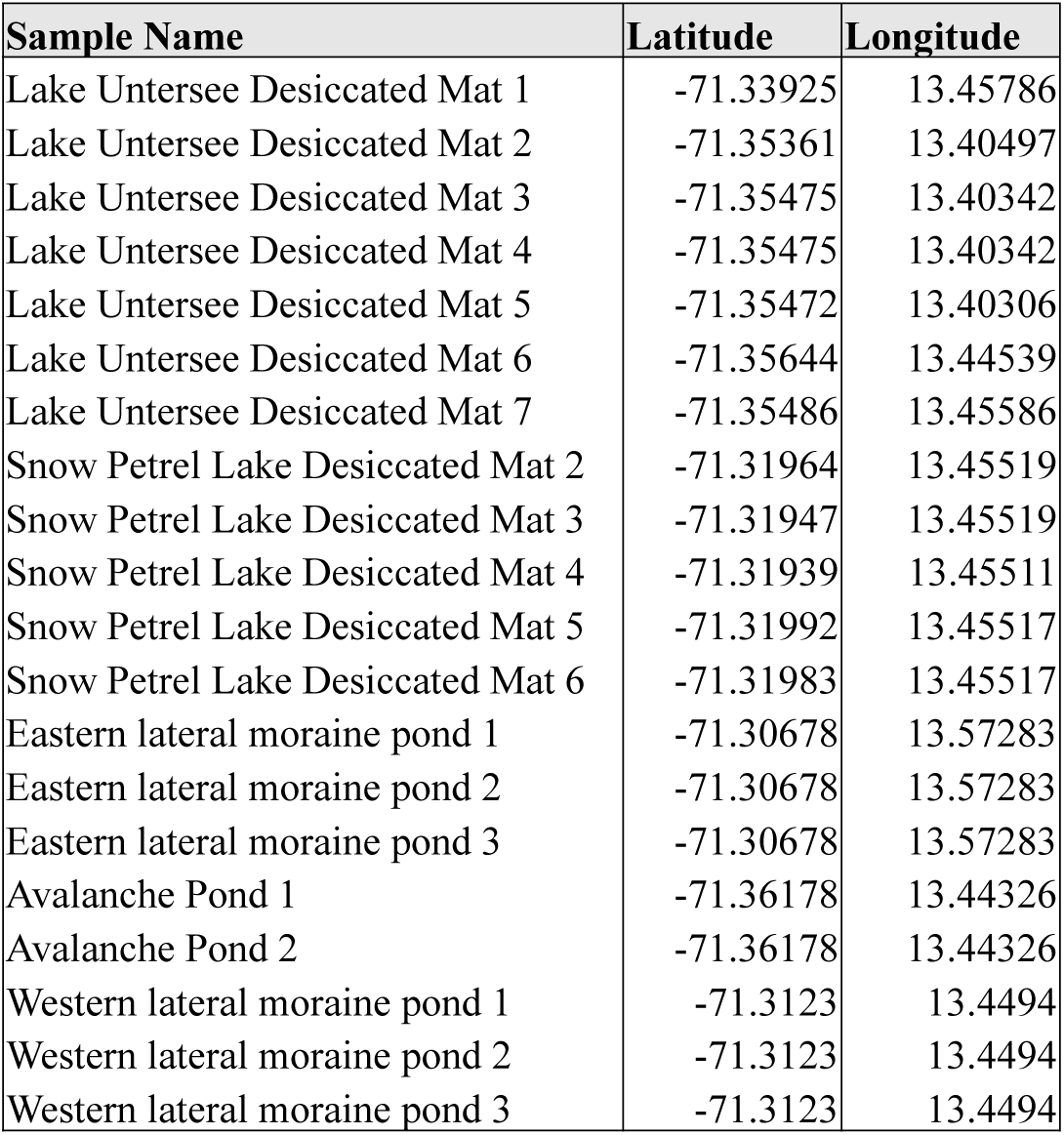
Sampling coordinates for microbial mat collections from the Lake Untersee region, Antarctica. Geographic coordinates (decimal degrees, WGS84) for desiccated microbial mats and pond communities sampled from Lake Untersee basin, Dronning Maud Land, East Antarctica. Replicate samples from identical coordinates represent biological replicates.

### DNA extraction and sequencing

Freeze-dried mats were subsequently used for DNA extraction using the QIAGEN DNeasy PowerBiofilm DNA Isolation Kit (MO BIO Laboratories, Carlsbad, CA, United States), in accordance with the manufacturer’s protocol. Bacterial 16S rRNA genes were amplified using the 515f–806r primers, and eukaryotic 18S rRNA genes were amplified using the 1391f–EukBr primer sets (Amaral-Zettler et al. 2009; Caporaso et al. 2012). PCR products from both marker genes were cleaned using the AxyPrep Mag PCR Clean Up Kit (Axygen, New York, NY, United States), quantified with a Qubit fluorometer (Invitrogen, United Kingdom) before being pooled together. Sequencing was performed on the Illumina MiSeq platform at the Natural History Museum, London. Sequences generated in this study (pond and desiccated mats) are deposited under BioProject PRJNA1366640, while previously published benthic mat sequences are available under PRJNA638357; both datasets were jointly analyzed in this manuscript.

### Sequence processing, ASV identification, and diversity analysis

Sequenced data were processed by first demultiplexing the data using idemp (https://github.com/yhwu/idemp) and trimming primers with Cutadapt (Martin 2011). Quality plots were visually inspected, and sequences were processed using the DADA2 (Callahan et al. 2016) pipeline to infer Amplicon Sequence Variants (ASVs) differing by at least one nucleotide. Filtering parameters were: 16S rRNA gene reads trimmed to 250/230 nucleotides (forward/reverse) and 18S rRNA gene reads trimmed to 115/110 nucleotides, both with maximum expected errors of 5 per read direction. All sequences were truncated at quality score 2 with removal of sequences containing ambiguous nucleotides (truncQ=2, maxN=0). Paired-end reads were merged, and chimeras and singletons were removed.

Taxonomic assignment was performed against the SILVA database (silva_nr_v138_train_set.fasta) for 16S rRNA gene and the PR2 database (pr2_version_4.11.1_dada2.fasta) for 18S rRNA gene sequences. The “mctoolsr” R package was used to filter out chloroplasts, mitochondria, and eukaryotic reads from the 16S rRNA gene data and bacterial reads from the 18S rRNA gene data (Leff 2017). ASV tables were rarified to the number of sequences in the lowest populated sample (20,804 for 16S rRNA and 1812 for 18S rRNA gene respectively). Alpha diversity metrics (ASVs richness, Faith’s phylogenetic diversity (Faith’s PD; (Faith 1992) and Pielou’s Evenness) were calculated from the rarefied community matrices.

We calculated beta diversity using the weighted UniFrac distance metric (Lozupone and Knight 2005) implemented in the “phyloseq” R package (McMurdie and Holmes 2013), and Bray–Curtis dissimilarity implemented in the “mctoolsr” R package (Leff 2017). Principal Coordinates Analysis (PCoA) ordinations were computed on UniFrac dissimilarities to illustrate differences in the phylogenetic composition of microbial communities. To generate statistical support for these differences we ran the nonparametric PERMANOVA test (Anderson 2001) with 999 permutations with the “adonis” function in the “vegan” R package (Oksanen et al. 2019). To evaluate whether samples differed in their dispersion, an analysis of multivariate homogeneity (PERMDISP; (Anderson 2006) was run using the function “betadisper” with the “vegan” R package using default parameters.

Taxonomic composition differences among habitats (lake, pond, and desiccated) at phylum, order, family, and genus levels were assessed using Kruskal-Wallis tests for each taxon, followed by pairwise Wilcoxon tests with Bonferroni correction for post-hoc comparisons. False discovery rate (FDR) adjustment was applied across all taxa within each taxonomic level to account for multiple testing.

Indicator species analysis for ASV indicators of lake, pond and desiccated habitats was performed with the “multipatt” function (with the “rg” association function) in the “indicspecies” R package (Cáceres and Legendre 2009) using 999 permutations for significance testing. This analysis was conducted separately for bacterial (16S rRNA gene) and eukaryotic (18S rRNA gene) communities, as well as for cyanobacteria alone within the 16S rRNA gene dataset, to identify taxa that showed significant associations with specific habitat types. For the full bacterial dataset, only taxa with high indicator values (≥0.875) were retained to focus on the strongest environmental indicators given the large number of ASVs. For cyanobacterial and eukaryotic analyses, a more inclusive threshold of ≥0.7 was applied due to the smaller number of ASVs in these datasets, ensuring sufficient indicators were retained for meaningful ecological interpretation while still maintaining robust habitat-specific associations.

### Core microbiome analysis

Core microbiome analysis was performed to identify taxa consistently present across the three habitat types (lake benthic mats, pond mats and desiccated mats) using the “microbiome” R package (Lahti et al., 2017). Rarefied count data were converted to phyloseq objects and transformed to compositional abundances. Core microbiome heatmaps were generated to visualize the prevalence of individual ASVs across logarithmically spaced detection thresholds ranging from 0.001% to 95% relative abundance. Detection thresholds were logarithmically spaced to provide fine resolution at low abundances where many taxa transition from present to absent, while maintaining broader intervals at higher abundances where fewer taxa are affected. Taxa were ranked by average prevalence and displayed with genus-level taxonomic identification when available, with higher taxonomic levels used as fallbacks for unclassified taxa. Core taxa were defined as those present at >0.1% relative abundance in >90% of samples within each dataset. This analysis was conducted separately for bacterial (16S rRNA gene) and eukaryotic (18S rRNA gene) communities to identify the most persistent and abundant taxa across the combined environments.

### Beta nearest taxon index (βNTI) analysis

β nearest taxon index analysis was performed to assess the relative contributions of deterministic versus stochastic processes in bacterial and eukaryotic community assembly across pond, lake, and desiccated mat environments. Rarefied 16S and 18S rRNA gene sequence data were matched with a phylogenetic tree constructed from representative sequences, and phylogenetic distances were calculated using the cophenetic distance method from the ape package in R. βNTI values were computed with 1000 randomizations, 4 parallel workers, and weighted abundance data. Pairwise βNTI comparisons were categorized as within-environment or between-environment comparisons. Values of | βNTI| > 2 indicated dominance of deterministic processes (heterogeneous selection for βNTI > +2, homogeneous selection for βNTI < -2), while | βNTI| ≤ 2 indicated stochastic processes. Statistical differences in βNTI values among environments were tested using analysis of variance (ANOVA) followed by Tukey’s honest significant difference test for pairwise comparisons. All analyses were conducted using the “iCAMP”, “picante”, and “ape” R packages.

### Spatial patterns and distance-decay relationships

To assess spatial patterns in bacterial community structure, we performed Mantel tests to examine the relationship between geographic distance and community dissimilarity (Legendre and Legendre 2012). Only pond and desiccated mat samples were included in this analysis, as lake mat samples did not have coordinates associated with them. Geographic distances between sample pairs were calculated using the Haversine formula to account for Earth’s curvature (Hijmans 2017). Community dissimilarity matrices were computed using Bray-Curtis dissimilarity based on rarefied ASV abundance data (20,804 sequences per sample for 16S and 1812 for 18S rRNA gene). Mantel tests were performed using Pearson correlation with 999 permutations to assess the significance of distance-decay relationships (Mantel 1967). We conducted three separate analyses: (1) all samples combined to assess landscape-scale patterns, (2) pond samples only, and (3) desiccated mat samples only, to test for habitat-specific dispersal limitations (Martiny et al. 2006). All analyses were performed in R using the “vegan” package (Oksanen et al., 2019). Significant positive correlations between geographic and community distance matrices indicate distance-decay relationships consistent with dispersal limitation (Green et al. 2004; Martiny et al. 2011).

### Source Tracer analysis

To investigate source environments for microbial communities in this study, we used SourceTracker2, a Bayesian community-wide culture-independent microbial source tracking algorithm (Knights et al. 2011). Source-sink relationships among pond, lake, and desiccated habitats were analyzed by conducting three separate analyses, each designating one habitat as sink and the other two as potential sources. SourceTracker2 was run using the Gibbs sampling algorithm on non-rarefied ASV count data to estimate the proportional contribution of each source habitat to sink sample microbial communities.

### Network analysis

Co-occurrence networks were constructed separately for bacterial (16S rRNA gene) and eukaryotic (18S rRNA gene) communities, as well as a combined cross-domain network using the “mctoolsr” and “igraph” R packages. Prior to network construction, both 16S and 18S rRNA gene ASV tables were filtered to remove contaminant and unclassified taxa and underwent initial global filtering, removing features below a threshold (0.025 for 16S and 0.022 for 18S rRNA gene respectively) to screen out globally rare taxa and account for differences in sequencing depth between bacterial and eukaryotic datasets. Taxa were further filtered to retain only those present in at least 20% of the samples within each environment, with counts greater than 0.01% relative abundance before being reduced to the 100 most abundant ASVs per domain per habitat for the final analysis. To address the compositional nature of the data, the final filtered ASV counts underwent a Centered Log-Ratio (CLR) transformation after a pseudocount of 0.5 was added to all counts using the “compositions” R package. Co-occurrence was determined by calculating Spearman rank correlation (ρ) coefficients using the “Hmisc” R package between ASVs, and only strong associations (∣ρ∣≥0.8) were retained as network edges. Networks were visualized in both single-domain and combined 16S-18S rRNA gene formats, where nodes (ASVs) were sized by degree and edges differentiated by sign (positive/negative correlation) and combined networks utilized a custom double-ring layout.

### Functional gene analysis

To assess stress adaptation strategies and functional capacity across habitats, we conducted PICRUSt2 analysis (Phylogenetic Investigation of Communities by Reconstruction of Unobserved States) to predict metagenomic content from 16S rRNA gene sequences (Douglas et al. 2020). PICRUST2 infers functional gene content by matching amplicon sequence variants to phylogenetically related reference genomes and produces predicted abundances for 7,166 KEGG Orthology (KO) groups across all samples. From the complete PICRUST2 output, we extracted predicted abundances for 63 stress-related genes of interest by matching their KEGG Orthology identifiers to the predicted KO profiles. Stress-related genes were manually curated into six functional categories based on their documented roles in environmental stress responses: General Stress (genes involved in multiple, non-specific stress pathways such as chaperones and general protection mechanisms), Oxidative Stress (genes specifically involved in reactive oxygen species management and redox homeostasis), Osmotic/Salt (genes involved in osmotic balance and salt tolerance), Starvation/Stringent (genes involved in nutrient limitation responses and stringent control), Heat_Cold_Shock (genes involved in thermal adaptation and freeze tolerance), and DNA Repair (genes involved in maintaining genome integrity under stress conditions) (Supplementary Table 1). Genes with documented functions spanning two or more of these stress categories were classified as Multi-Stress genes, reflecting their pleiotropic roles in stress adaptation. Additionally, we extracted 24 CRISPR-associated (Cas) genes representing components of Type I, II, III, V, and VI CRISPR systems, with each Cas gene predicted independently based on its KEGG Orthology identifier (Supplementary Table 2). Gene abundances for both stress and CRISPR-Cas gene sets were normalized by dividing each gene’s predicted abundance by the total predicted gene content per sample, yielding relative abundances that account for differences in sequencing depth and total functional capacity across samples. For stress genes, we selected based on their relevance to extreme Antarctic conditions and microbial survival strategies in oligotrophic environments (Cary et al. 2010; Ramasamy et al. 2023), resulting in seven final categories with 5-12 genes each (mean = 9, coefficient of variation = 29.4%). Compositional differences and relative abundances of stress gene categories were compared among the three habitat types (lake mats, pond mats, and desiccated mats) using multivariate statistical approaches including PERMANOVA, SIMPER analysis, and non-metric multidimensional scaling (NMDS) to identify environment-specific functional signatures and stress adaptation patterns. SIMPER analysis identified stress gene categories contributing most to pairwise dissimilarity between habitats, with size-corrected contributions calculated as per-gene effect sizes (dissimilarity/gene count) and enrichment ratios (observed/expected contribution; >1.5 = over-represented, <0.67 = under-represented). Indicator species analysis (multipatt) with FDR correction identified habitat-specific stress gene categories. The same statistical framework was applied to analyze CRISPR gene composition and abundance patterns across environments.

In addition, functional guilds were predicted from bacterial 16S rRNA gene sequences using FAPROTAX v1.2.4 (Louca et al. 2016), which assigns ecological and metabolic functions based on literature-derived annotations. Functional abundance data were normalized, and only categories with a relative abundance ≥ 0.0005 across samples were retained. Analyses focused on the top 15 most abundant functions, including nitrogen, sulfur, and methane cycling, and organic matter degradation. Functional community structure was assessed using Principal Coordinates Analysis (PCoA) based on Bray-Curtis dissimilarities, and statistical differences between habitats (lake, pond and desiccated mats) were tested with Kruskal-Wallis tests and post-hoc comparisons. Functional diversity metrics, including richness and evenness were calculated to evaluate variation across habitats. Habitat-specific metabolic signatures were visualized using horizontal bar plots. Because many Antarctic lineages are poorly represented in reference databases, functional profiles inferred from 16S rRNA gene sequences using PICRUSt2 and FAPROTAX need to be interpreted cautiously.

## Results

### Microbial community composition and distribution across lake, ephemeral ponds and desiccated mat habitats

We assessed microbial community composition across three distinct habitats, including lake benthic mats, ephemeral ponds, and desiccated mats, to examine how bacterial and eukaryotic assemblages vary with environmental context. Bacterial community composition differed significantly among habitats at all investigated taxonomic levels (PERMANOVA: all p < 0.05). At the phylum level, all mat types were dominated by Cyanobacteria (36-47%) but showed distinct secondary patterns (Fig. 2A). Lake communities exhibited higher Gammaproteobacteria and notably heterogeneous composition, with tufts from the top of cone shaped mats differing markedly from other lake samples. Pond environments were characterized by elevated Bacteroidia (p < 0.001) and Chloroflexia (p < 0.001). Desiccated mats showed more diverse phylum-level composition with notable Actinobacteria (p = 0.001), Alphaproteobacteria, and Acidimicrobia. Order-level analysis revealed finer taxonomic structuring, with lake communities distinguished by high Cyanobacteriales (p < 0.001), while ponds showed elevated Oxyphotobacteria_Incertae_Sedis (p < 0.001), RD011 (p < 0.001), and Chloroflexales (p = 0.001), and desiccated environments exhibited diverse orders including Micrococcales, Flavobacteriales, and Leptolyngbyales. At the family level, Phormidiaceae dominated lake mats (p < 0.001), Chloroflexaceae distinguished pond communities (p = 0.001), and desiccated mats showed high diversity with Microbacteriaceae, Flavobacteriaceae, Leptolyngbyaceae, and Caulobacteraceae. Genus-level analysis identified *Tychonema* CCAP 1459-11B as the dominant lake taxon, with *Leptolyngbya* ANT.L67.1 and *Chloronema* characterizing ponds, and desiccated mats exhibiting the highest diversity including *Tychonema* CCAP 1459-11B, *Leptolyngbya* ANT.L67.1, *Flavobacterium*, *Brevundimonas*, and *Cryobacterium*. Taxonomic composition at order, family, and genus levels are shown in Supplementary Fig. S1 (A, B, C), with exact percentages and statistical results provided in Supplementary Tables. These taxonomic patterns demonstrate consistent environment-specific community structuring across all phylogenetic levels examined.

**Figure 2.**
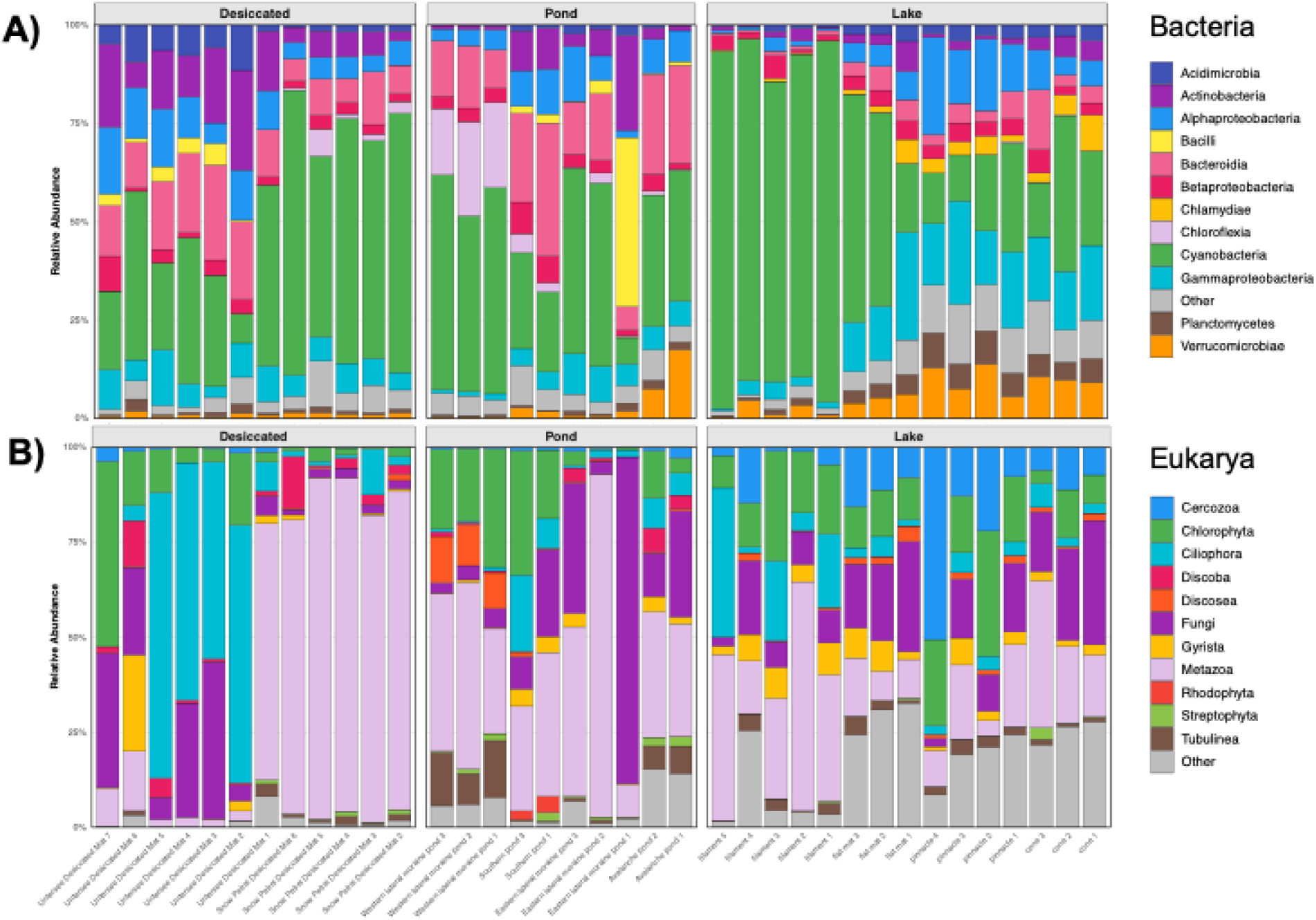
Bacterial and Eukaryotic community composition across habitats. Relative abundance of major bacterial (A) and eukaryotic (B) phyla in desiccated terrestrial mats, pond, and lake communities. Each bar represents a sample. Taxonomic assignments were performed using the SILVA reference database for 16S rRNA sequences and the PR2 reference database for 18S rRNA sequences.

Eukaryotic community composition differed significantly among habitats at all taxonomic levels (PERMANOVA: all p < 0.05), with significant differences in within-group dispersion among environments (betadisper: p < 0.05), indicating that compositional differences reflected both mean shifts and variable community structuring within habitats. In contrast to bacterial communities dominated by a single phylum, eukaryotic assemblages exhibited much higher taxonomic diversity with no clear dominant taxon (Fig. 1B). At the phylum level, communities showed distinct environmental-specific characteristics with desiccated environments characterized by high microfauna (rotifers, tardigrades, nematodes), Ciliophora, and Fungi, reflecting the greater complexity of eukaryotic life strategies. Pond communities showed elevated microfauna, Fungi, and Chlorophyta, while lake environments were distinguished by higher Cercozoa (p < 0.001), Chlorophyta, and microfauna. At finer taxonomic resolution, environment-specific assemblages emerged clearly. Lake communities were dominated by the rotifer *Adineta vaga*, green algae *Chlorococcum*, and nematodes (Plectidae sp.; p < 0.001), alongside diverse protistan grazers including Cercozoa and ciliates. Pond communities supported specialized extremophile assemblages including the amoeba *Hartmannella vermiformis* (p = 0.009 vs. lakes), psychrophilic yeasts (Sporobolomyces; p = 0.031 vs. lakes), and tardigrades (*Acutuncus antarcticus*). Desiccated mats showed highest microfaunal diversity with abundant rotifers (*Adineta vaga*), the psychrophilic yeast *Glaciozyma antarctica* (p = 0.003 vs. ponds), Hypotrichia ciliates, and diverse cold-adapted protists. Taxonomic composition at order, family, and genus levels are shown in Supplementary Fig. S2 (A, B, C), with exact percentages and statistical results provided in Supplementary Tables. These taxonomic patterns demonstrate consistent habitat-specific community structuring across all phylogenetic levels examined.

### Environment-specific taxa identified through indicator species and core microbiomes

Indicator species analysis identified strong environment-specific ASV associations with distinct phylum-level patterns. Lake bacterial indicators were dominated by Verrucomicrobiae (Verrucomicrobiaceae, *Terrimicrobium*) reflecting elevated lake Verrucomicrobiae abundance (6.3 ± 4.3%), alongside diverse Chlamydiae and Planctomycetes (Fig. 3A). Pond indicators consisted primarily of Bacteroidia (Cytophagales, *Hymenobacter*) consistent with elevated pond Bacteroidia (18.2 ± 8.4%), as well as Chloroflexi, Gammaproteobacteria (*Leptothrix*), and cyanobacterial taxa (*Phormidium*). Desiccated environments showed fewer indicators, predominantly Bacteroidia and Actinobacteria taxa. Cyanobacterial analysis revealed complete habitat specialization, with lake-specific *Tychonema,* pond-associated *Phormidium*, and desiccated *Leptolyngbya* and *Chamaesiphon* taxa. Although Cyanobacteria were abundant across all environments (36–47%), indicator ASVs were entirely environment-specific, highlighting strain-level niche partitioning within this dominant photosynthetic lineage (Fig. 3B). Eukaryotic indicators displayed a balanced distribution across habitats. Lake communities were distinguished by diverse protists (such as Cercozoa, Frontoniidae, and Oxytricha), nematodes (Plectidae), the alga *Nannochloropsis limnetica*, and by several fungal taxa which, due to limitations in reference databases, could only be identified to higher taxonomic levels (e.g., *Fungi*, *Ascomycota*). Pond environments were characterized by the amoeba *Hartmannella vermiformis*, the flagellate *Tetramitia* and micrometazoans (notably rotifers). In contrast, desiccated mats were marked by the chytrid *Hyaloraphidium curvatum*, various protists (Oxytrichidae, Vahlkampfiidae), and *Adineta* sp. rotifers. (Fig. 3C).

**Figure 3.**
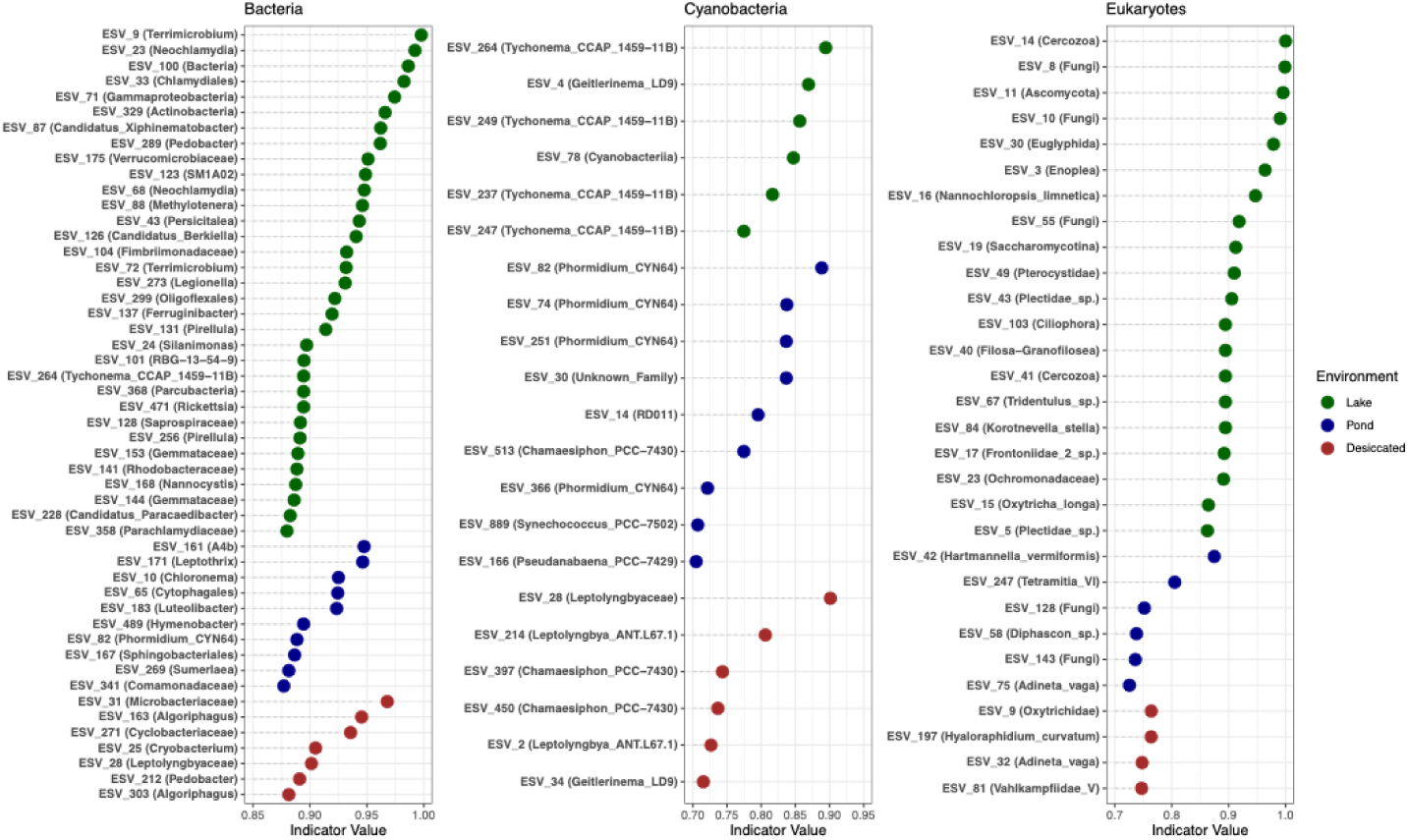
Environment-specific indicator species across microbial communities. Indicator species analysis showing ASVs with significant habitat associations for bacterial, cyanobacterial, and eukaryotic communities across lake, pond, and desiccated environments. Points represent indicator values (strength of environmental association), with higher values indicating stronger habitat specificity.

Core microbiome analysis revealed distinct patterns between bacterial (16S rRNA gene) (Fig. 4A) and eukaryotic (18S rRNA gene) (Fig. 4B) communities across the combined pond, lake, and desiccated mat habitats. At the 0.1% detection threshold, bacterial taxa showed moderate prevalence with 38 taxa (2.6% of all bacterial ASVs) present in ≥50% of samples, though only 2 taxa achieved ≥90% prevalence and just 1 taxon was universally present (100% prevalence). The most prevalent bacterial taxa included the heterotrophic bacterium *Galbitalea*, followed by *Rhodanobacter*, *Ilumatobacter*), and members of the Planctomycetes (*Pirellula*). The core bacterial community contracted rapidly as detection thresholds increased, with only 4 taxa maintaining ≥50% prevalence at 1% abundance and a single taxon persisting at 10% abundance. In contrast, the eukaryotic community demonstrated a more constrained but potentially more stable core, with 14 taxa (3.4% of all eukaryotic ASVs) present in ≥50% of samples at 0.1% detection threshold and 2 taxa achieving universal presence. The eukaryotic core was dominated by bdelloid rotifers (*Adineta vaga*), green algae (*Chlorococcum microstigmatum*, Chlamydomonadales), ciliated protists (*Oxytricha longa*), and fungi. *Adineta vaga* remained highly prevalent across intermediate thresholds. Notably, eukaryotic taxa showed greater resilience at intermediate abundance levels, with 3 taxa maintaining ≥50% prevalence and 1 taxon achieving ≥90% prevalence at the 1% detection threshold, though no taxa persisted at 10% abundance.

**Figure 4.**
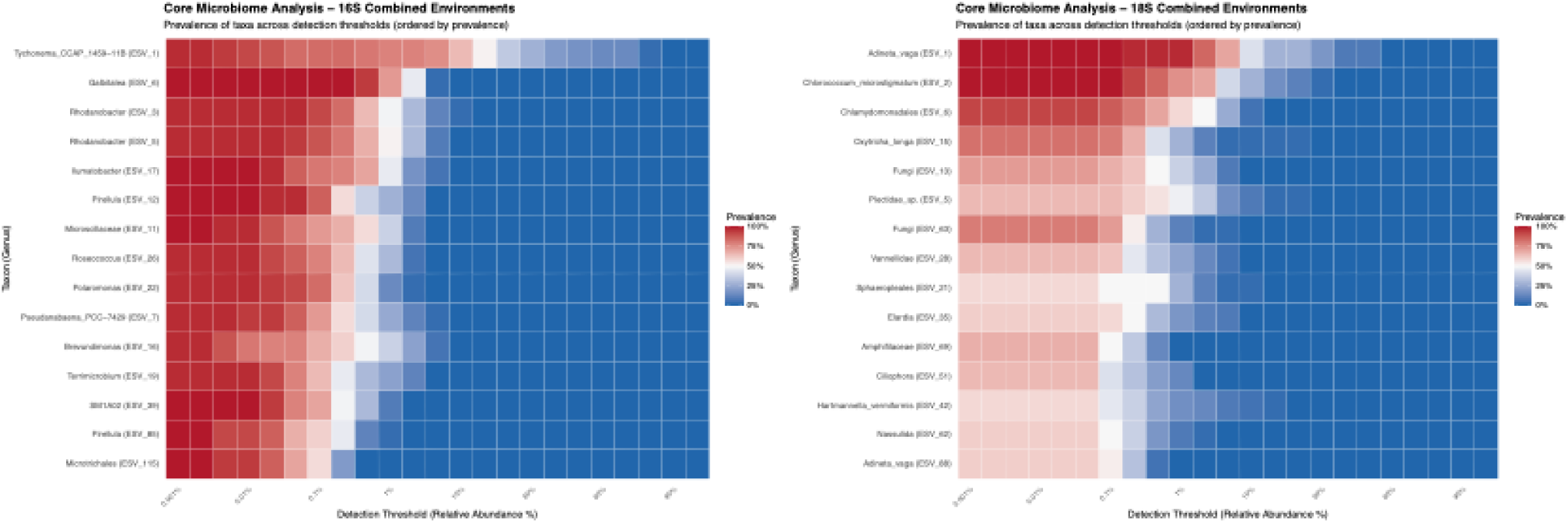
Core microbiome analysis reveals taxa prevalence across detection thresholds. Heatmaps showing the prevalence of (left) bacterial (16S rRNA) and (right) eukaryotic (18S rRNA) taxa across different relative abundance detection thresholds. Taxa are ordered by overall prevalence, with red indicating high prevalence (present in most samples) and blue indicating low prevalence (present in few samples) at each abundance threshold. ASV numbers indicate exact sequence variants for each taxon.

### Assembly processes: deterministic versus stochastic structuring

β-nearest taxon index (βNTI) analysis revealed that community assembly processes varied significantly between bacterial (Fig. 5A) and eukaryotic communities (Fig. 5B) and across the three habitat types. Overall, both domains demonstrated predominantly stochastic assembly across environments, with significant variation in βNTI values reflecting habitat-specific trends. Bacterial communities showed moderate but significant environmental differentiation in assembly processes (ANOVA: F₂,₂₁₃ = 5.22, p = 0.006), with lake and pond habitats both differing significantly from desiccated habitats (Tukey HSD: p = 0.018 and p = 0.014, respectively), while pond and lake bacterial assembly processes were statistically similar (p = 0.79). Mean within-environment βNTI values indicated stochastic assembly in all bacterial habitats (desiccated = -1.27, lake = -0.84, pond = -0.73), with desiccated environments showing the strongest trend toward deterministic processes and the highest variability. In contrast, eukaryotic communities displayed much stronger environmental structuring of assembly mechanisms (ANOVA: F₂,₂₁₃ = 55.31, p < 0.001), with lake communities differing markedly from both desiccated and pond habitats (Tukey HSD: p < 0.001 for both comparisons), while desiccated and pond eukaryotic assembly processes were similar (p = 0.78). Mean βNTI values showed that eukaryotic assembly was also predominantly stochastic, with lake communities near-neutral (βNTI = 0.037) and desiccated and pond communities showing moderate trends toward homogeneous selection (desiccated = -1.03, pond = -0.93), reflecting slight environmental filtering in ephemeral habitats without reaching deterministic thresholds (|βNTI| > 2).

**Figure 5.**
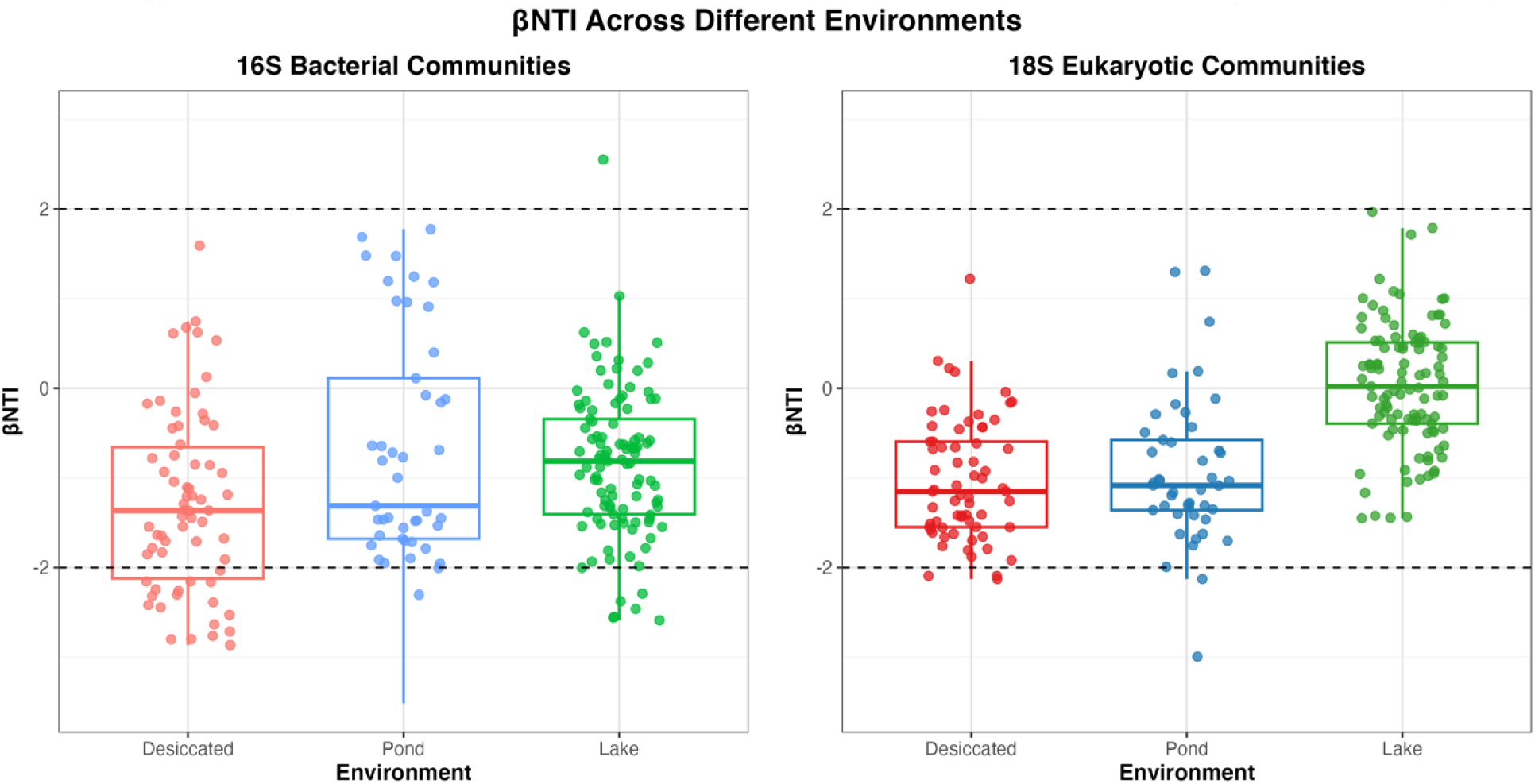
Community assembly processes across environments. βNTI values for bacterial (16S rRNA gene) (left panel) and eukaryotic (18S rRNA gene) (right panel) communities from desiccated, pond, and lake habitats. Dashed lines indicate deterministic assembly thresholds (±2).

### Spatial structure and distance-decay relationships

Spatial analysis of bacterial and eukaryotic communities revealed habitat-specific distance-decay relationships, with desiccated mats exhibiting the strongest spatial structuring for both prokaryotes and eukaryotes. Bacterial communities demonstrated significant distance-decay relationships within individual habitat types (Fig. 6A). The combined analysis across all environments showed a significant positive relationship (Mantel r = 0.316, p = 0.002), though within-habitat comparisons (Mat-Mat, Pond-Pond) displayed positive distance-decay while cross-habitat comparisons (Mat-Pond) showed a negative relationship. Desiccated mats displayed the strongest bacterial spatial pattern (Mantel r = 0.70, p = 0.001), while pond communities also showed a strong and significant relationship (Mantel r = 0.58, p = 0.001), despite operating across a relatively small spatial scale (9.08 km maximum distance). Eukaryotic communities showed even stronger distance-decay relationships in both habitats, with desiccated mats displaying the strongest pattern (Mantel r = 0.84, p = 0.001) compared to ponds (Mantel r = 0.56, p = 0.001). (Fig. 6B).

**Figure 6.**
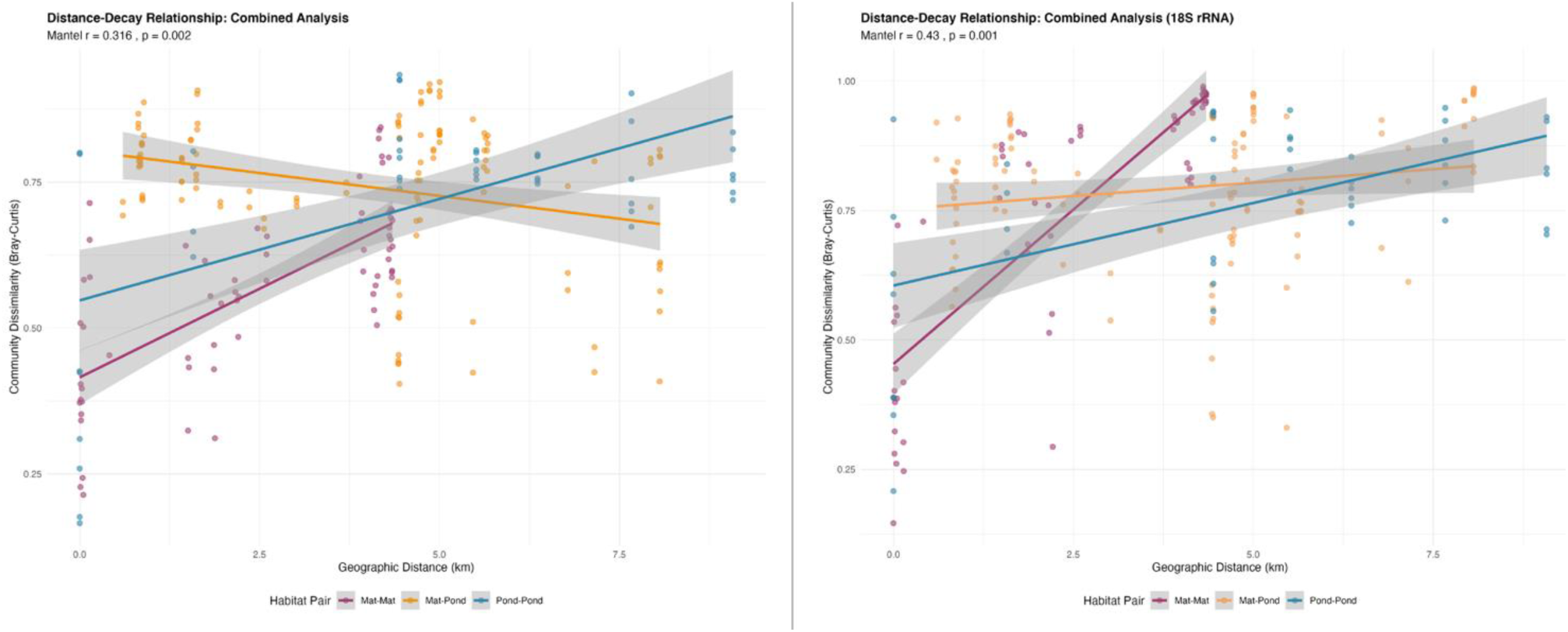
Habitat-specific distance-decay patterns in bacterial and eukaryotic communities. Community dissimilarity (Bray-Curtis) plotted against geographic distance for (left panel) bacterial (16S rRNA) and (right panel) eukaryotic (18S rRNA) communities across habitats. Each point represents a pairwise comparison between samples, colored by habitat pair type. Trend lines show fitted relationships with 95% confidence intervals of the three habitat pair categories: Mat-Mat (Pairwise comparison between two samples from desiccated mat habitats); Mat-Pond (Pairwise comparison between a sample from a desiccated mat habitat and a sample from a pond habitat); Pond-Pond (Pairwise comparison between two samples from pond habitats)

### Source-sink dynamics and dispersal connectivity

Source tracking analysis revealed contrasting dispersal patterns between prokaryotic and eukaryotic communities across the three habitat types. Prokaryotic communities showed substantial cross-habitat exchange, with desiccated mats serving as the primary source environment (Fig. 7A). Lake communities derived 60.8% of their microbiome from desiccated sources with minimal pond contribution (0.7%) and because desiccated mats originate from lake benthic mats, this apparent contribution largely reflects shared origin rather than an independent dispersal flux back into the lake. Pond communities received 28.4% from desiccated mats and only 1.8% from under-ice lake benthic communities. Desiccated mat communities displayed the most balanced exchange pattern, receiving approximately equal contributions from both lake (30.3%) and pond (31.5%) sources. In contrast, eukaryotic communities exhibited minimal cross-habitat dispersal (Fig. 7B), with much of each community (83.7-98%) attributed to unknown sources rather than the other sampled environments. The limited eukaryotic exchange was primarily between pond and desiccated environments, with pond communities receiving 15.2% from desiccated sources, while lake-pond and lake-desiccated exchanges remained below 4% in all cases.

**Figure 7.**
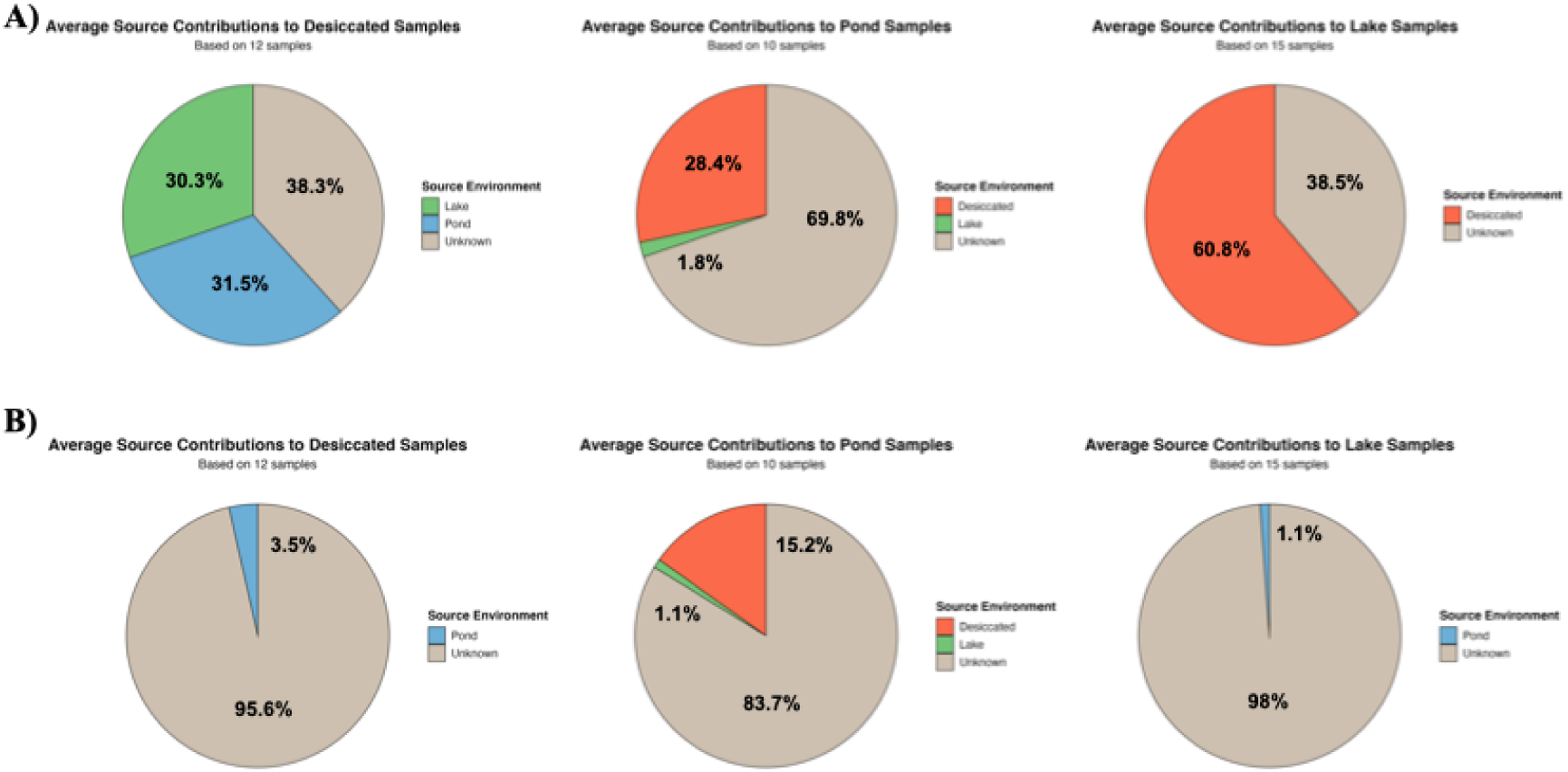
Source-sink dynamics across lake, pond, and desiccated mat habitats. SourceTracker2 analysis showing average proportional contributions of source environments to sink communities for (A) prokaryotic (16S rRNA) and (B) eukaryotic (18S rRNA) communities. Each pie chart represents the mean source contributions across all samples for each sink environment (Lake: n=15 samples; Pond: n=10 samples; Desiccated: n=12 samples). Colored segments indicate proportional contributions from identified source habitats (Desiccated: red; Lake: green; Pond: blue), while gray segments represent unknown sources not attributable to the sampled habitats.

### Cross-domain association networks and assembly

Microbial co-occurrence network analysis revealed a strong gradient of complexity (Fig. 8), with the lake habitat exhibiting the lowest network density (249 total edges, 41 cross-domain edges), suggesting minimal overall co-occurrence. In contrast, the pond (1,059 edges) and desiccated (967 edges) mat habitats were highly interconnected. The primary distinction lay in the nature of their inter-domain relationships (16S-18S rRNA gene): pond mats featured the highest absolute number of cross-domain edges (377), which were predominantly positive (65.0%), indicating mutualistic or shared habitat preference. Conversely, the highly dense desiccated network, while featuring 295 cross-domain edges, was uniquely dominated by negative correlations (53.6%), suggesting that competitive or exclusionary associations govern the microbial structure in this dry habitat.

**Figure 8.**
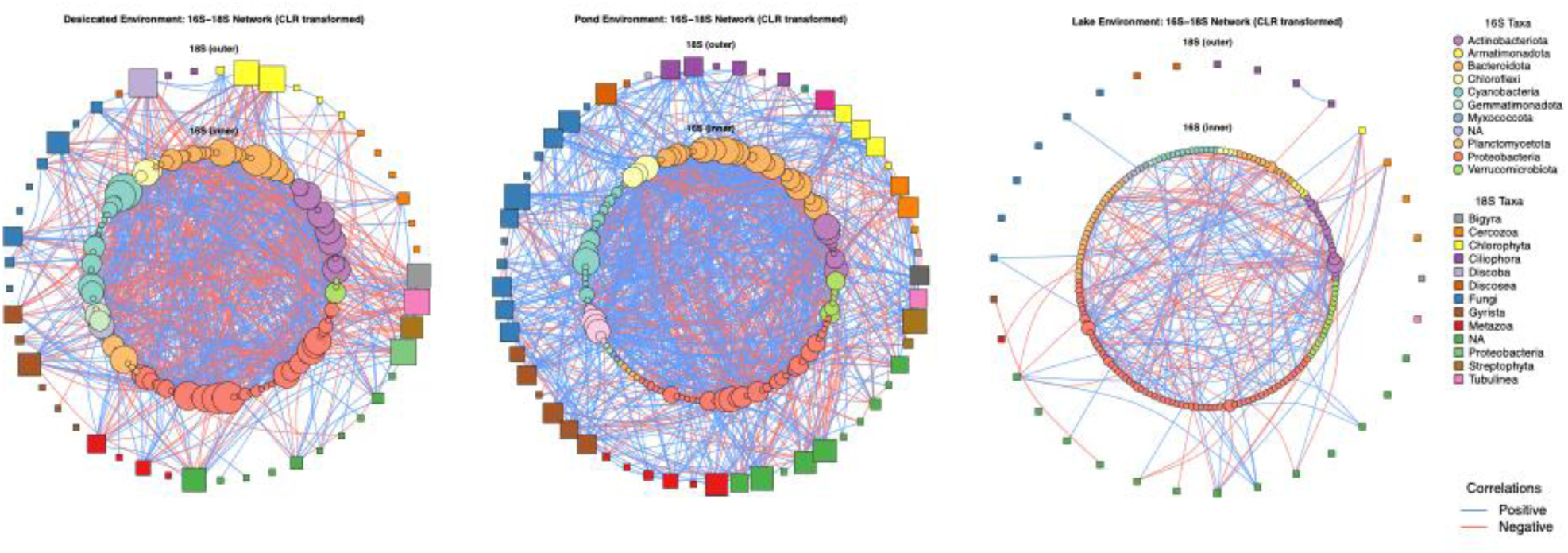
Microbial co-occurrence networks across lake, pond, and desiccated mat habitats. Co-occurrence networks of prokaryotic (16S rRNA gene, outer ring) and eukaryotic (18S rRNA gene, inner ring) communities in lake, pond, and desiccated mat habitats constructed using CLR-transformed abundance data. Node colors represent different taxonomic groups as indicated in the legends. Edge colors represent correlation types: blue edges indicate positive correlations, red edges indicate negative correlations.

### Functional trait differentiation across habitats

FAPROTAX analysis revealed distinct functional profiles across the three study habitats. Principal coordinates analysis (PCoA) demonstrated clear separation of functional communities, with the first two axes explaining 85.8% of the total variance (PC1: 75.7%, PC2: 10.1%; Figure 9A). Lake communities clustered separately from both desiccated mat and pond communities, while desiccated and pond samples showed intermediate similarity. Photosynthesis-related processes dominated the functional landscape across all habitats. The 15 most abundant functional categories included phototrophy, oxygenic photoautotrophy, photoautotrophy, and photosynthetic cyanobacteria, which collectively comprised the majority of predicted functional abundance (Figure 9B). Despite this overall dominance of photosynthetic functions, several other categories exhibited significant enrichment in certain habitat types. Ureolysis showed significant enrichment in pond samples (p < 0.001), while intracellular parasites were most abundant in lake habitats (p < 0.001). Biogeochemical cycling functions related to nitrogen, sulfur, carbon, and methane metabolism represented a smaller but ecologically critical component of the predicted functional repertoire (Figure 9C). Lake habitats showed significant enrichment in multiple nitrogen cycling processes, including nitrogen fixation (p < 0.05), nitrification (p < 0.001), and aerobic ammonia oxidation (p < 0.01). Lake habitats also exhibited significantly higher methylotrophic activity, with both methylotrophy (p < 0.001) and methanol oxidation (p < 0.001) being most abundant in these permanently hydrated habitats. In contrast, desiccated mat communities showed selective enrichment in chitinolysis (p < 0.05), suggesting presence of specialized organic matter degradation capabilities under water-limited conditions. Lake communities exhibited the highest functional richness, while pond communities showed intermediate richness and desiccated mat communities displayed the lowest richness (Figure 9D). However, this difference was not statistically significant across the habitats (Kruskal-Wallis p=0.05; Dunn’s post-hoc test P_adj_>0.05 for all comparisons). Functional evenness followed an opposite pattern, being highest in desiccated mats and lowest in lake samples.

**Figure 9.**
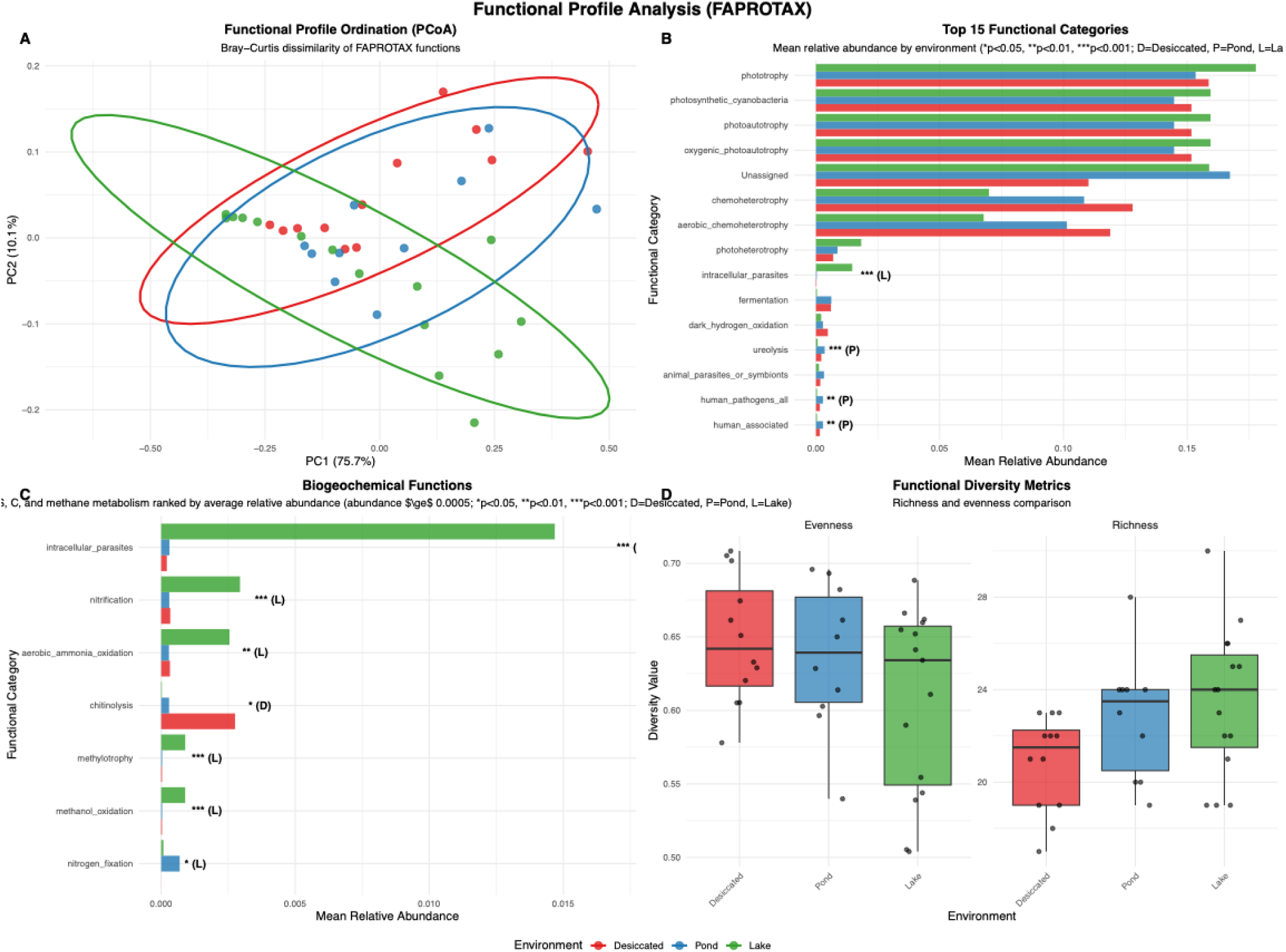
Predicted functional profile analysis reveals distinct metabolic specialization across environments. (A) Principal coordinates analysis (PCoA) of FAPROTAX-predicted functional profiles showing clear separation among desiccated mat (red), pond (blue), and lake (green) communities based on Bray-Curtis dissimilarity. (B) Mean relative abundance of the top 15 functional categories across environments. (C) Next most abundant biogeochemical cycling functions related to nitrogen, sulfur, carbon, and methane metabolism. (D) Functional diversity metrics comparing richness and evenness among environments. Significance levels: *p<0.05, **p<0.01, ***p<0.001; D=Desiccated, P=Pond, L=Lake indicate which environment shows significant enrichment.

PICRUSt2 analysis of stress-related gene categories revealed significant functional differentiation among habitats, with clear separation evident in NMDS ordination (PERMANOVA: R² = 0.255, p = 0.003; NMDS stress = 0.062). The predicted stress gene repertoire showed consistent relative abundances across all habitats with normalized compositional data. SIMPER analysis identified Multi-Stress and Starvation_Stringent response categories as the primary drivers of community differentiation, with Multi-Stress showing the highest enrichment levels (2.70-3.06x over-represented based on size-corrected analysis) and both Multi-Stress and Starvation_Stringent displaying consistently elevated size-corrected contributions (35.9% and 13.3%, respectively) across all pairwise comparisons, while Heat_Cold_Shock genes were consistently under-represented (0.23-0.45x enrichment). The strongest functional differentiation occurred between desiccated mat and lake mat habitats (90% dissimilarity), followed by desiccated mat versus pond mats (80% dissimilarity) and pond mat versus lake mat habitats (77% dissimilarity). Across all habitats, the dominant stress gene categories based on size-corrected contributions were Multi-Stress (35.9%), General_Stress (14.6%), Starvation_Stringent (13.3%), DNA_Repair (11.6%), and Osmotic_Salt (12.0%).

Interestingly, CRISPR-Cas gene composition also differed significantly among sample types (PERMANOVA: R² = 0.72, F = 7.89, p = 0.001), with pond habitats exhibiting the highest CRISPR gene richness followed by lake and desiccated mat communities. A total of 18 CRISPR genes were detected. While PICRUST2-based CRISPR predictions have inherent limitations due to the high variability of these systems, the relative patterns across habitat types appear robust. Within pond habitats, substantial variation in CRISPR abundance was observed among different geographic locations, while desiccated and lake communities exhibited more uniform CRISPR investment across samples. The top genes contributing to compositional differences included cas1, cas10, and cas2.

## Discussion

### Microbial community composition and habitat-specific adaptations

The significant bacterial community differentiation across pond, lake, and desiccated mat habitats at all taxonomic levels demonstrates strong environmental filtering consistent with extreme niche partitioning in Antarctic freshwater microbiomes . While Cyanobacteria universally dominated (36-47%) due to their exceptional adaptation to polar conditions including desiccation tolerance, UV resistance, and temperature flexibility (Martin-Andres et al. 2024), distinct secondary patterns emerged reflecting habitat-specific selective pressures. Lake Untersee’s benthic communities inhabit a chronically carbon-depleted, extremely oligotrophic, high-pH system with very low TIC/TOC and a sealed water column (Marsh et al. 2020), conditions that strongly constrain metabolic strategies and favor long-term trait specialization. Lake communities exhibited elevated Gammaproteobacteria alongside Cyanobacteria, suggesting tight phototroph-heterotroph coupling in stable, light-replete conditions. The identification of habitat-specific cyanobacterial strains (*Leptolyngbya* ANT.L67.1, *Tychonema* CCAP 1459-11B) supports evidence for cold adapted Antarctic cyanobacteria (Lumian et al. 2024b). The elevated abundance of Actinobacteria, particularly the Micrococcales order and *Cryobacterium* genus, in desiccated mats and pond habitats, reflects their specialized adaptations to fluctuating water availability and oligotrophic conditions. This distribution pattern aligns with the ecological preferences of these taxa, as *Cryobacterium* species represent typical Antarctic psychrotrophic bacteria whose genomes encode extensive stress response mechanisms, including genes for oxidative stress tolerance, cold-stress adaptation, and general environmental stress resistance (Teoh et al. 2024). The preference of Actinobacteria for water-stressed environments is further supported by studies demonstrating their dominance in arid Antarctic Dry Valley soils (18-52% of communities), with abundances significantly decreasing during experimental wetting events, confirming their adaptation to desiccated, oligotrophic conditions rather than permanently hydrated habitats (Niederberger et al. 2019). This diversity of aquatic and terrestrial habitats likely supports microbial survival and recolonization under extreme Antarctic conditions, maintaining both active and dormant populations capable of responding to environmental fluctuations (Stone et al. 2024; Juarez Rivera et al. 2025).

The distinct eukaryotic community patterns observed across environments reflect the greater ecological complexity and environmental sensitivity of eukaryotic life strategies compared to bacterial communities. Molecular studies of Antarctic eukaryotic diversity have revealed much higher taxonomic complexity than previously suspected, with fungi, algae, and protists showing distinct environment-specific distribution patterns (Lawley et al. 2004). The betadispersion analysis (PERMANOVA, p < 0.05) revealing significant within-habitat variability suggests that microhabitat heterogeneity and microscale environmental gradients play a stronger role in structuring eukaryotic than bacterial communities. Cercozoa were notably abundant in lake habitats but nearly absent from pond and desiccated mat communities. As primarily bacterivorous or omnivorous protists, cercozoans are typically abundant in environments with stable moisture and sustained microbial prey availability (Fiore-Donno et al. 2019). Fungal community composition also showed striking habitat-specific patterns. Basidiomycota were detected almost exclusively in desiccated mats and pond habitats, while lake environments harbored only Ascomycota fungi. This distribution aligns with the general exclusion of Basidiomycota from permanently aquatic habitats, where Ascomycota typically dominate (Shearer et al. 2007). The presence of Basidiomycota in desiccated mats and ponds, but not lakes, suggests these taxa colonize during dry phases or from terrestrial sources, thriving in environments with intermittent water availability that more closely resemble their typical soil and litter habitats. The psychrophilic basidiomycete *Glaciozyma antarctica*, detected predominantly in desiccated samples, exemplifies this pattern as a species originally isolated from sea ice and terrestrial moss rather than permanently submerged habitats (Turchetti et al. 2011). *Glaciozyma antarctica* possesses extensive genomic adaptations for sustained responses to temperature variations and environmental stress, suggesting it thrives in habitats with fluctuating rather than stable conditions (Firdaus-Raih et al. 2018). Metazoan distribution patterns further differentiated the three habitat types. Tardigrada were present only in select desiccated and pond samples but nearly absent from lake environments, while nematodes displayed the opposite pattern with higher abundance in lakes compared to ponds and desiccated mats. This differential distribution reflects distinct ecological strategies: tardigrades are quintessentially adapted to semi-terrestrial environments experiencing periodic desiccation, where their exceptional anhydrobiotic capabilities provide a selective advantage (Tsujimoto et al. 2016). In permanently submerged lake habitats, tardigrades’ cryptobiosis adaptations offer no competitive benefit and may represent metabolic costs that favor obligately aquatic nematodes. Antarctic nematodes such as Plectidae species, though capable of anhydrobiosis (Adhikari et al. 2009), were notably more abundant in permanently wet lake habitats where continuous moisture permits sustained bacterial grazing. Collectively, these habitat-specific eukaryotic assemblages demonstrate that environmental stability, water permanence, and desiccation frequency act as primary filters structuring Antarctic microbial mat communities at multiple trophic levels.

### Indicator Species analysis reveals fine-scale environmental specialization beyond community structure patterns

While relative abundance analysis demonstrated broad environmental filtering patterns, indicator species analysis revealed the specific taxonomic drivers of habitat differentiation and uncovered fine-scale ecological specialization. Indicator ASVs represent taxa that exhibit strong statistical associations with particular environmental conditions, serving as reliable biological indicators of habitat-specific selective pressures. Lake bacterial indicators were dominated by Verrucomicrobia, Chlamydiae, and Planctomycetes. Verrucomicrobia are recognized as important polysaccharide degraders in freshwater lakes, utilizing algal and cyanobacterial exudates through diverse glycoside hydrolases (Martinez-Garcia et al. 2012; Chiang et al. 2018). Some of the highest pond indicator ASVs were affiliated with Chloroflexi, consistent with their roles as metabolically versatile heterotrophs and photoheterotrophs common in microbial mat communities (Klatt et al. 2013; Mehrshad et al. 2018). Beyond these high-ranking Chloroflexi ASVs, pond habitats demonstrated strong selection for Bacteroidia indicators, particularly Cytophagales and *Hymenobacter*, consistent with their elevated overall Bacteroidia abundance. Recent studies have highlighted the ecological importance of these taxa in polar environments (Klassen and Foght 2011; Tanner et al. 2018; Terashima et al. 2019). The reduced number of indicators in desiccated mat habitats, dominated by Bacteroidia and Actinobacteria, reflects the harsh selective pressures that limit which taxa can serve as reliable indicators of extreme desiccation stress.

Most significantly, cyanobacterial indicator analysis revealed complete habitat specialization at the strain level despite uniform phylum-level dominance across all environments. This demonstrates that ecological niche partitioning operates at much finer taxonomic scales than revealed by abundance-based analyses alone (Jungblut et al. 2010; Tromas et al. 2018; Panwar et al. 2022). The environment-specific distribution of *Tychonema* (Lake), *Phormidium* (Ponds), and *Leptolyngbya* and *Chamaesiphon* (Desiccated mats) indicates that even within the dominant photosynthetic lineage, distinct evolutionary lineages have specialized for specific Antarctic microhabitat conditions. This pattern aligns with recent phylogenetic studies revealing extensive cryptic diversity within polar cyanobacterial genera and strong biogeographic patterns at the strain level (Lumian et al. 2024b; Durieu et al. 2025). Strain-level specialization supports the concept that apparent redundancy at higher taxonomic levels masks significant functional and ecological differentiation (Auladell et al. 2022). Antarctic studies further show that closely related strains can exhibit dramatically different stress tolerances, photosynthetic capabilities, and habitat preferences (Lumian et al. 2024a). The complete turnover of cyanobacterial indicator species between environments, despite consistent phylum-level dominance, reveals that environmental filtering in extreme systems can operate at fine taxonomic scales rather than broad functional categories. Such strain-level turnover and environment-specific functional differentiation mirror patterns observed in other Antarctic microbial communities (Vick-Majors et al. 2014) and suggest that while photosynthetic capability allows cyanobacteria to dominate all habitats, specific physiological adaptations determine competitive success at the strain level. These patterns highlight the importance of considering microbial diversity beyond broad phylogenetic or functional classifications when assessing ecosystem responses to environmental change.

Eukaryotic indicator analysis revealed pronounced habitat specificity, with most significant differences distinguishing lake habitats from pond and desiccated mats, reflecting the fundamental ecological divide between hydrated and desiccation-tolerant eukaryotic life strategies in these Antarctic systems. Lake habitats yielded the greatest number of eukaryotic indicators and were characterized by ciliate indicators (Frontoniidae, *Oxytricha*) and the microalga *Nannochloropsis limnetica*, taxa that rely on stable aquatic conditions for their active growth and reproduction. Pond and desiccated mat habitats yielded fewer indicators with lower values. Notably, the bdelloid rotifer *Adineta vaga* emerged as an indicator of both pond and desiccated mat environments but not lakes, reflecting the genus’s renowned capacity for anhydrobiosis through constitutively expressed antioxidants and DNA repair pathways (Hespeels et al. 2023). Pond habitats were further characterized by the cosmopolitan bacterivorous amoeba *Hartmannella vermiformis*, a species widely distributed in freshwater habitats that commonly serves as a host for bacterial endosymbionts (Delafont et al. 2018). Additionally, the flagellate *Tetramitia* and rotifers as pond indicators suggest these habitats support active bacterivory and filter-feeding strategies typical of shallow productive waters. The chytrid *Hyaloraphidium curvatum* represents an unusual non-flagellated aquatic fungus whose presence in desiccated mats may reflect tolerance of intermittent hydration. These patterns reinforce the primary ecological divide between permanently aquatic and intermittently hydrated habitats as the dominant driver of eukaryotic community structure.

### Domain-divergent core microbiomes: bacterial colonization breadth versus eukaryotic taxonomic persistence

While indicator species analysis identified habitat specialists, core microbiome analysis reveals taxa that transcend habitat boundaries and persist across environmental gradients. Our results suggest that while bacterial communities harbor greater overall diversity, eukaryotic communities maintain a more consistently present core assemblage across the three habitat types. The bacterial community’s moderate prevalence at low detection thresholds but rapid contraction at higher abundances suggests a strategy of widespread occurrence at low abundances, with few taxa achieving consistent dominance across habitats. Conversely, the eukaryotic community’s smaller but more persistent core, particularly the universal presence of the bdelloid rotifer *Adineta vaga*, reflects the known ecological dominance of bdelloids in Antarctic terrestrial and aquatic environments (Wada et al. 2023). The persistence of *A. vaga* across all habitat types suggests this species functions as an ecologically significant grazer that maintains ecosystem connectivity through active dispersal between aquatic and terrestrial environments. The higher resilience of eukaryotic taxa at intermediate abundance thresholds, despite their overall lower diversity, supports the hypothesis that larger organisms like bdelloid rotifers experience greater dispersal limitation but achieve more stable population dynamics once established, contributing to the habitat-specific assembly patterns observed in our βNTI analysis.

### Stochastic assembly dominates both prokaryotic and eukaryotic communities despite stronger environmental sensitivity in eukaryotes

The predominance of stochastic assembly processes across both bacterial and eukaryotic communities, despite strong environmental differentiation in composition, suggests that stochastic dispersal establishes initial community composition before environmental filtering drives habitat-specific differentiation. While mean within-environment βNTI values consistently fell within the stochastic range (-2 to +2), eukaryotic communities exhibited significantly stronger environment-specific assembly patterns than bacteria (F = 55.31 vs. 5.22). This pattern aligns with Antarctic freshwater lake studies where stochastic processes accounted for up to 95% of eukaryotic community assembly despite clear environmental structuring (Zhang et al. 2022) likely reflecting the overriding influence of dispersal limitation and ecological drift in isolated microhabitats (Vyverman et al. 2010; Logares et al. 2018).The negative βNTI values in terrestrial environments suggest moderate trends toward homogeneous selection, wherein environmental filtering favors phylogenetically similar lineages adapted to shared stress conditions (Stegen et al. 2013; Ning et al. 2020). While not reaching deterministic thresholds (|βNTI| > 2), these values indicate that desiccation stress, freeze-thaw cycles, and UV exposure weakly favor related taxa with convergent stress tolerance traits (Goberna et al. 2014; Dini-Andreote et al. 2015).

Lake eukaryotic communities exhibited near-neutral processes (βNTI = 0.037), suggesting stable aquatic conditions reduce selective pressures and allow stochastic processes to dominate more completely (Stegen et al. 2012; Aguilar and Sommaruga 2020). However, Lake Untersee’s long-term “stability” is punctuated by rare but substantial glacial lake outburst floods (GLOFs) that periodically raise lake level, alter water chemistry, and deliver carbon subsidies on Holocene timescales (Faucher et al. 2021b). The stronger environmental structuring of eukaryotic assembly processes reflects interactions between body size, dispersal limitation, and niche breadth. Larger-bodied eukaryotes experience greater dispersal constraints than bacteria (Logares et al. 2018; Luan et al. 2020), making them more susceptible to local environmental filtering even when stochastic processes dominate (Li et al. 2023). This dispersal limitation amplifies the importance of local environmental conditions, creating stronger environment-specific assembly patterns even within a fundamentally stochastic framework. Our findings challenge the traditional view that extreme environments should be dominated by deterministic assembly through strong environmental filtering. Instead, they support emerging perspectives suggesting that extreme environments can promote stochastic assembly when harsh conditions limit both regional diversity and dispersal connectivity (Lemoine et al. 2023). The Antarctic landscape around Lake Untersee exemplifies this scenario: geographic isolation, limited connectivity among habitats, and relatively small microbial populations create conditions where ecological drift and dispersal limitation prevent deterministic selection from fully structuring communities (Vyverman et al. 2010).

### Significant habitat-specific spatial patterns with different responses in bacteria and eukaryotes

Our analysis revealed significant distance-decay relationships in both bacterial and eukaryotic communities, with patterns varying substantially between habitat types but showing consistent trends across microbial groups (bacterial Mantel r = 0.316-0.70; eukaryotic Mantel r = 0.43-0.84 indicating moderate to strong distance-decay). These findings demonstrate that geographic distance influences microbial community assembly even at relatively small spatial scales, consistent with growing evidence for spatial structuring in Antarctic microbial communities (Bottos et al. 2020).

Both bacterial and eukaryotic communities exhibited consistently stronger spatial structuring in desiccated mats compared to ponds, suggesting shared ecological constraints across domains. Bacterial communities exhibited stronger spatial structuring in desiccated mats (Mantel r = 0.70) compared to ponds (Mantel r = 0.58), likely reflecting the more isolated nature of mat environments and limited dispersal opportunities during desiccation. This pattern contrasts with the regional connectivity observed in other Antarctic pond systems, where bacterial communities showed extensive sharing across locations up to 40 km apart (Archer et al. 2015). The stronger spatial isolation in our desiccated mats suggests these communities may represent locally adapted populations that persist in place during periods of environmental stress. Eukaryotic communities displayed the same pattern, with even stronger spatial structuring in desiccated mats (Mantel r = 0.84) compared to ponds (Mantel r = 0.56). The strong distance-decay in mat eukaryotes may reflect the larger size and more complex life cycles of eukaryotic organisms, which could make dispersal more challenging in desiccated terrestrial environments where liquid water is only transiently available. The consistent pattern across both prokaryotic and eukaryotic domains suggests that the physical isolation of desiccated mat environments and constraints on dispersal during desiccation represent fundamental barriers to microbial movement regardless of organism size or complexity.

The detection of significant spatial patterns at small scales (maximum 9.08 km for both habitats) indicates that local processes likely contribute strongly to community assembly in these extreme environments. This aligns with studies from the Antarctic Dry Valleys, where significant distance-decay relationships were observed but became weaker after accounting for environmental variation, suggesting that both dispersal limitation and environmental filtering contribute to spatial patterns (Bottos et al. 2020). Our findings are consistent with broader patterns in microbial biogeography, where the strength of distance-decay relationships has been shown to vary between different environments and habitats (Clark et al. 2021). The significant combined analyses for both bacteria (r = 0.316, p = 0.002) and eukaryotes (r = 0.43, p = 0.001) indicate that geographic distance influences community composition across the entire landscape. However, the negative distance-decay relationship observed in cross-habitat comparisons of Bacteria (Mat-Pond pairs) suggests that local environmental filtering between habitat types is stronger than geographic distance effects, with nearby desiccated mats and ponds being more dissimilar than distant ones of the same habitat type. This supports findings from other Antarctic freshwater systems where environmental factors dominated over dispersal in driving community structure, despite high physical connectivity between habitats (Ramoneda et al. 2021). The habitat-specific nature of our distance-decay relationships suggests that desiccated mats and ephemeral ponds function as distinct ecological islands with limited cross-habitat dispersal, potentially maintaining unique microbial assemblages that contribute to regional diversity around Lake Untersee.

### Desiccated mats show unexpected prokaryotic community mixing, contrasting with habitat-specific eukaryotic communities

Source-tracking analysis revealed striking domain-specific dispersal patterns, though interpretation requires careful consideration of methodological limitations inherent to SourceTracker when applied to physically connected habitats. In this system, SourceTracker should be interpreted primarily as revealing overlap in community composition rather than true directional dispersal, because the apparent high contribution of desiccated mats to lake prokaryotic communities (60.8%) likely represents a methodological artifact rather than true ecological dispersal, as desiccated mats physically originate from benthic lake environments through buoyancy-driven detachment. Thus, the apparent contribution of desiccated mats to lake communities reflects shared ancestry and circular source–sink definitions rather than an ecologically meaningful flux, because this self-referential relationship, where the “source” is actually derived from the “sink”, creates spurious connectivity in SourceTracker analyses that cannot distinguish true dispersal from shared ancestry (Knights et al. 2011). Previous work demonstrated that cryoconite communities from the Anuchin glacier represent the primary external source for lake Untersee benthic mats, contributing approximately 36% of community composition through meltwater inputs and subglacial discharge (Weisleitner et al. 2019). This glacier-derived input, rather than pond or desiccated mat dispersal, likely represents the dominant external source shaping lake microbial communities.

When accounting for this methodological limitation, the more robust ecological patterns emerge from comparisons not confounded by circular source–sink relationships, such as between pond and desiccated mat communities. Desiccated mats displayed balanced prokaryotic contributions from both lake (30.3%) and pond (31.5%) sources. This pattern suggests that during the extended period desiccated mat fragments spend on the ice surface after emerging from the lake benthos, they experience colonization by wind-dispersed prokaryotes from surrounding terrestrial environments. The mechanism likely involves aeolian transport of microorganisms from exposed pond surfaces and soils onto desiccated mat surfaces, where they establish alongside the original lake-derived community (Scott et al. 2025). Similarly, the 28.4% contribution from desiccated mats to pond communities represents biologically meaningful dispersal, potentially occurring when desiccated fragments are deposited into ponds through wind transport or when pond formation occurs in areas where desiccated materials have accumulated. These patterns suggest that desiccated mats are not merely passive reservoirs of diversity; rather, they maintain a mixture of locally adapted and dispersal-derived taxa that could become metabolically active under favorable conditions, functioning as both an active community and a repository of ecological potential. Few studies have compared microbial contributions across lakes, ponds, and desiccated mats within the same Antarctic catchment, highlighting the novelty of our multi-habitat source-tracking approach. The minimal direct lake-pond prokaryotic connectivity (0.7-1.8%) reveals that despite occupying the same catchment, these aquatic habitats function as largely isolated systems with limited hydrological exchange. This isolation likely reflects the absence of permanent surface water connections between Lake Untersee and surrounding ponds, with ephemeral meltwater streams providing insufficient dispersal opportunities to establish significant microbial exchange (Šabacká et al. 2012). Wind-mediated transport of desiccated mat fragments and associated prokaryotic communities may function as the main mechanism connecting otherwise isolated aquatic habitats in this hydrologically constrained Antarctic system (Pearce et al. 2009; Bottos et al. 2020).

In striking contrast to substantial prokaryotic exchange, eukaryotic communities exhibited minimal cross-habitat dispersal, with the vast majority (83.7-98.0%) attributed to unknown sources rather than the sampled environments. This pattern aligns with findings from Antarctic freshwater lakes where stochastic processes accounted for up to 95% of microbial eukaryotic community assembly, driven by severe dispersal limitation in isolated aquatic systems (Zhang et al. 2022). The limited eukaryotic exchange detected was primarily between pond and desiccated environments (15.2% from desiccated to pond), with virtually no detectable contributions involving lake communities (<4% in all cases). This domain-specific contrast reflects fundamental differences in dispersal capacity: eukaryotic microorganisms, being generally larger-bodied with more complex life cycles and often requiring specific environmental cues for dormancy and reactivation, experience greater constraints on passive aerial dispersal compared to smaller, more resilient prokaryotic cells (Wilkinson et al. 2012; Ptacnik et al. 2010). Additionally, many Antarctic eukaryotes may rely on dispersal vectors operating at regional or continental scales, such as bird-mediated transport or long-distance atmospheric circulation, explaining why the majority of eukaryotic community composition could not be attributed to local source environments within the immediate Lake Untersee catchment (Vyverman et al. 2010; Logares et al. 2018).

These contrasting dispersal patterns have important implications for understanding ecosystem connectivity and resilience in Antarctic landscapes. While prokaryotic communities maintain connectivity across terrestrial habitats through wind dispersal, receiving external inputs primarily from glacier-derived cryoconite for lake environments and atmospheric deposition for ponds and desiccated mats, eukaryotic communities appear functionally isolated at local scales (Nemergut et al. 2013; Convey et al. 2014). Collectively, these patterns suggest that desiccated mats may also act as inocula for more extreme, arid, or frozen environments, consistent with relic Antarctic mat studies showing long-term preservation of metabolic potential, lipids, and genomic structure during extended inactivity (Zaikova et al. 2019; Lezcano et al. 2022; Drozd et al. 2023). Even during low-activity phases, desiccated mats may therefore function simultaneously as dormant archives and as vectors facilitating microbial exchange across the broader landscape.

### From sparse aquatic to dense competitive networks: water availability drives microbial-eukaryotic associations

The sparse network structure observed in lake habitats suggests that stable, permanently hydrated conditions promote independent niche occupation with minimal inter-species dependencies. The low cross-domain connectivity in lakes indicates that bacterial and eukaryotic communities can function relatively independently when environmental conditions are favorable and predictable. In contrast, the highly interconnected networks in pond and desiccated environments reflect the increased importance of biotic interactions under environmental variability and stress. The predominance of positive cross-domain correlations in pond environments (65.0% positive) provides support for the stress gradient hypothesis under moderate environmental stress (He et al. 2013). Our results show co-occurrence patterns consistent with shifts from more positive to more negative associations along stress gradients, while acknowledging that correlation networks cannot distinguish shared habitat preference from mutualism or competition. Seasonal pond habitats, characterized by fluctuating water availability and resource inputs, appear to select for mutualistic relationships between bacteria and eukaryotes that enhance survival during stress periods. These positive associations likely reflect complementary resource utilization, cross-feeding relationships, or shared habitat preferences that buffer against environmental variability. The unique dominance of negative correlations in desiccated mat networks (53.6% negative) represents a striking shift toward competitive and exclusionary associations under extreme environmental stress. This pattern contradicts predictions of the stress gradient hypothesis for the most extreme environments, where facilitation might be expected to dominate. Instead, our results suggest that under extreme desiccation stress, resource limitation becomes so severe that competitive exclusion overrides potential benefits of cooperation. The prevalence of negative bacterial-eukaryotic associations in desiccated mat habitats may reflect fundamental differences in stress tolerance strategies between these domains, leading to incompatible resource requirements or antagonistic associations during the few periods when liquid water becomes available. The contrasting patterns of cross-domain co-occurrence patterns across habitats have important implications for ecosystem stability and resilience. Pond habitats, with their facilitative bacterial-eukaryotic networks, may exhibit greater functional redundancy and resistance to perturbations through cooperative buffering mechanisms. Conversely, the competitive networks in desiccated environments may be more vulnerable to cascading effects from species loss, as negative interactions can amplify disturbances throughout the community (Coyte et al. 2015). The intermediate complexity of desiccated networks, despite their competitive nature, may reflect a balance between the ecological necessity of maintaining some cooperative relationships while engaging in intense competition for limited resources. This suggests that even under extreme stress, complete competitive exclusion may be constrained by the requirement for minimal ecosystem functioning (Hernandez et al. 2021).

### Habitat-specific functional specialization drives ecosystem service differentiation in microbial communities

The clear functional differentiation among environments (85.8% total variance explained) suggests that habitat-specific selective pressures drive not only taxonomic but also functional community assembly (Zhang et al. 2025). This finding extends beyond simple abundance patterns to reveal how environmental filtering shapes the metabolic potential and ecosystem service provisioning of microbial communities across Antarctic microhabitats. While photosynthesis-related functions dominated all habitats, reflecting the fundamental importance of primary production in Antarctic ecosystems (Quesada and Vincent 2012), the observed functional specialization reveals that apparent metabolic redundancy masks significant niche partitioning. The universal prevalence of oxygenic photoautotrophy confirms cyanobacteria as the primary producers across Antarctic freshwater habitats, consistent with their well-documented evolutionary adaptation to polar environments (Chrismas et al. 2015). The significant enrichment of nitrogen cycling functions in lake habitats reflects the enhanced biogeochemical processing capacity of permanently hydrated habitats. The selective enrichment of chitinolysis in desiccated mat communities represents an adaptive response to the unique substrate availability and stress conditions of water-limited environments. Chitin degradation capabilities provide access to nitrogen and carbon from fungal cell walls and invertebrate exoskeletons, which may accumulate in desiccated environments due to reduced decomposition rates (Beier and Bertilsson 2013). Together, this functional specialization demonstrates how microbial communities adapt their metabolic repertoires to exploit available resources under extreme environmental conditions, supporting the concept of resource-driven functional selection in oligotrophic ecosystems.

### Stress gene functional analysis reveals environment-specific adaptation strategies despite conserved total stress capacity

The consistent total stress gene abundance across ehabitats (∼0.037-0.038 relative abundance) demonstrates that all Antarctic microbial communities maintain substantial investment in stress tolerance mechanisms, reflecting the fundamental importance of stress adaptation in polar ecosystems (Casanueva et al. 2010; Margesin and Miteva 2011). However, the significant compositional differences reveal habitat-specific optimization strategies, with Multi-Stress genes (35.9% size-corrected contribution, 2.70-3.06x enrichment) dominating community differentiation across all environments. The prevalence of Multi-Stress genes, which function across multiple stress pathways, alongside elevated Starvation_Stringent responses (13.3% contribution, 1.33-1.91x enrichment) indicates that Antarctic microbial communities rely on coordinated, multi-pathway stress responses rather than single-gene solutions to cope with complex and unpredictable environmental stressors (Varin et al. 2012). Conversely, Heat_Cold_Shock genes showed consistent under-representation (0.23-0.45x), suggesting that thermal adaptation, while present, is not the primary selective pressure in these perennially cold environments.

The convergent functional strategies between desiccated and pond habitats (67% dissimilarity) support the hypothesis that resource limitation drives similar adaptive responses regardless of water availability, reflecting shared selective pressures for efficient resource utilization and stress tolerance (Ács et al. 2019; George et al. 2023). Conversely, the unexpectedly high functional dissimilarity between pond and lake habitats (90%) despite both being aquatic suggests that water regime stability, rather than simple aquatic versus terrestrial classification, represents the primary selective pressure shaping stress adaptation strategies. The predominance of Multi-Stress genes and Starvation_Stringent responses, alongside moderate contributions from Oxidative_Stress tolerance and DNA_Repair, suggests these functions represent common core survival mechanisms in Antarctic habitats rather than cold-specific adaptations. This pattern reflects the fundamental physiological challenges in Antarctic terrestrial–aquatic environments, which are often characterized by ultra-oligotrophic conditions, variable conductivities, and high levels of oxygen production during the austral summer. The strong enrichment of pleiotropic Multi-Stress genes over specialized thermal responses highlights the importance of regulatory flexibility in unpredictable polar environments. The conserved total stress gene abundance across mats and other environments suggests that these communities harbor a ready-to-activate stress-response toolkit, supporting the view that desiccated mats function not just as preserved banks of diversity but as communities capable of rapid functional activation when conditions permit.

### Elevated CRISPR-Cas diversity in pond environments compared to lakes and desiccated mats suggests higher viral predation pressure

While PICRUST2-based predictions for highly variable systems like CRISPR-Cas have inherent limitations, the observed patterns across habitat types are ecologically consistent and statistically robust. A total of 18 CRISPR genes were detected, with pond environments showing the highest richness and diversity. The significantly higher CRISPR-Cas gene richness and diversity in pond ehabitats compared to desiccated and lake communities suggests that viral predation pressure varies substantially across the aquatic-terrestrial gradient. CRISPR-Cas systems function as adaptive immune mechanisms against viral and plasmid invasion, with community-level CRISPR diversity reflecting the historical diversity and intensity of phage-host interactions (Barrangou et al. 2007; Andersson and Banfield 2008). The elevated CRISPR-Cas Systems investment in ponds corresponds with their significantly higher network connectivity, suggesting that denser microbial interaction networks may facilitate more frequent viral transmission and horizontal gene transfer opportunities, though this is inferred rather than directly measured. These patterns are consistent with the known environmental instability of Antarctic pond systems, which experience strong fluctuations in water level, temperature, and chemistry (Faucher et al. 2021b), conditions that may intensify virus–host encounter rates. Network connectivity may directly amplify viral pressure by increasing contact rates between susceptible hosts and viral particles (Weitz et al. 2013), with higher connectivity potentially driving stronger selection for CRISPR-Cas defense systems. The 2-3 fold variation in CRISPR-Cas gene abundance among different pond locations further suggests that local environmental conditions modulating community interactions, such as nutrient availability, hydrological regime, or cell densities, drive corresponding variation in virus-host dynamics. In contrast, the reduced CRISPR-Cas system diversity in desiccated mat communities, despite their high overall network density, may reflect the predominance of competitive rather than facilitative interactions (53.6% negative correlations) that limit viral transmission efficiency. The sparse network structure in lake environments aligns with their intermediate CRISPR richness, suggesting that lower interaction frequencies constrain viral spread despite stable aquatic conditions that would otherwise support viral replication. Although these PICRUST2-derived patterns represent relative differences rather than absolute CRISPR abundances, the consistency across multiple Cas genes and alignment with known ecological drivers of CRISPR evolution (viral pressure, microbial density, environmental stability) support the biological relevance of these findings. Metagenomic validation would further strengthen conclusions about specific CRISPR system types and their functional roles across these habitats.

## Conclusions

Our study demonstrates that microbial communities across benthic lake mats, ephemeral ponds, and desiccated mats around Lake Untersee are shaped by a combination of strong environmental filtering, fine-scale taxonomic specialization, and divergent strategies between bacteria and eukaryotes. Bacterial communities exhibited broad dispersal, with few consistently dominant taxa, and complete strain-level turnover even within dominant cyanobacteria, while eukaryotes maintained smaller but more persistent cores constrained by strong dispersal limitation. Despite these differences, stochastic processes dominated assembly across habitats, with eukaryotes showing stronger habitat-specific structuring. Network analyses revealed facilitative interactions under moderate pond stress but competitive dominance in desiccated mats, underscoring how water availability drives cross-domain relationships. Functional predictions further indicated habitat-specific specialization, with lakes enriched in nitrogen and methane cycling functions, desiccated mats favoring stress tolerance and chitin degradation and ponds showing elevated CRISPR gene abundance, suggesting stronger viral selection pressure in these fluctuating environments. As these insights are derived from rRNA gene profiles, comprehensive metagenomic and metatranscriptomic studies will be essential to disentangle microbial connectivity, ecological interactions, and functional activity in greater detail, thereby refining our understanding of resilience and ecosystem services across Antarctic aquatic–terrestrial transitions. Future time-lapse metatranscriptomic experiments, spanning wetting, drying, and freeze–thaw transitions, will be particularly valuable for determining whether these communities actively remodel their metabolic responses or instead function largely as preserved reservoirs of dormant functional potential. Taken together, these aquatic–terrestrial transitions offer a natural terrestrial model for understanding how microbial communities assemble and persist in CO₂-limited, ice-covered ecosystems, providing insights relevant to analogous habitats proposed for Enceladus and early Mars.

## Author contributions

Lara Vimercati: Conceptualization; methodology; investigation; visualization; writing – original draft.

Imogen Chakrabarti: Investigation; writing – review and editing.

Annabel Lindley: Investigation; writing – review and editing.

Carla Greco: Investigation; writing – review and editing.

Dale T. Andersen: Supervision; funding acquisition; writing – review and editing.

Anne D. Jungblut: Conceptualization; supervision; funding acquisition; writing – review and editing.

## Acknowledgements

This work was supported by the TAWANI Foundation, the Trottier Family Foundation, NASA’s Exobiology program (Andersen: 80NSSC18K1094), and the Marie Skłodowska-Curie Actions Postdoctoral Fellowship (Vimercati: EP/X024873/1). Logistical support was provided by Ultima Antarctic Logistics, Cape Town, South Africa. We are grateful to COL (IL) Jennifer N. Pritzker, IL ARNG (Retired), President and Founder, TAWANI Foundation; Lorne Trottier of the Trottier Family Foundation and fellow field team members for their support during the expedition.

## Data availability statement

The Illumina reads generated in this study are openly available in the Sequence Read Archive with the accession code PRJNA1366640.

---

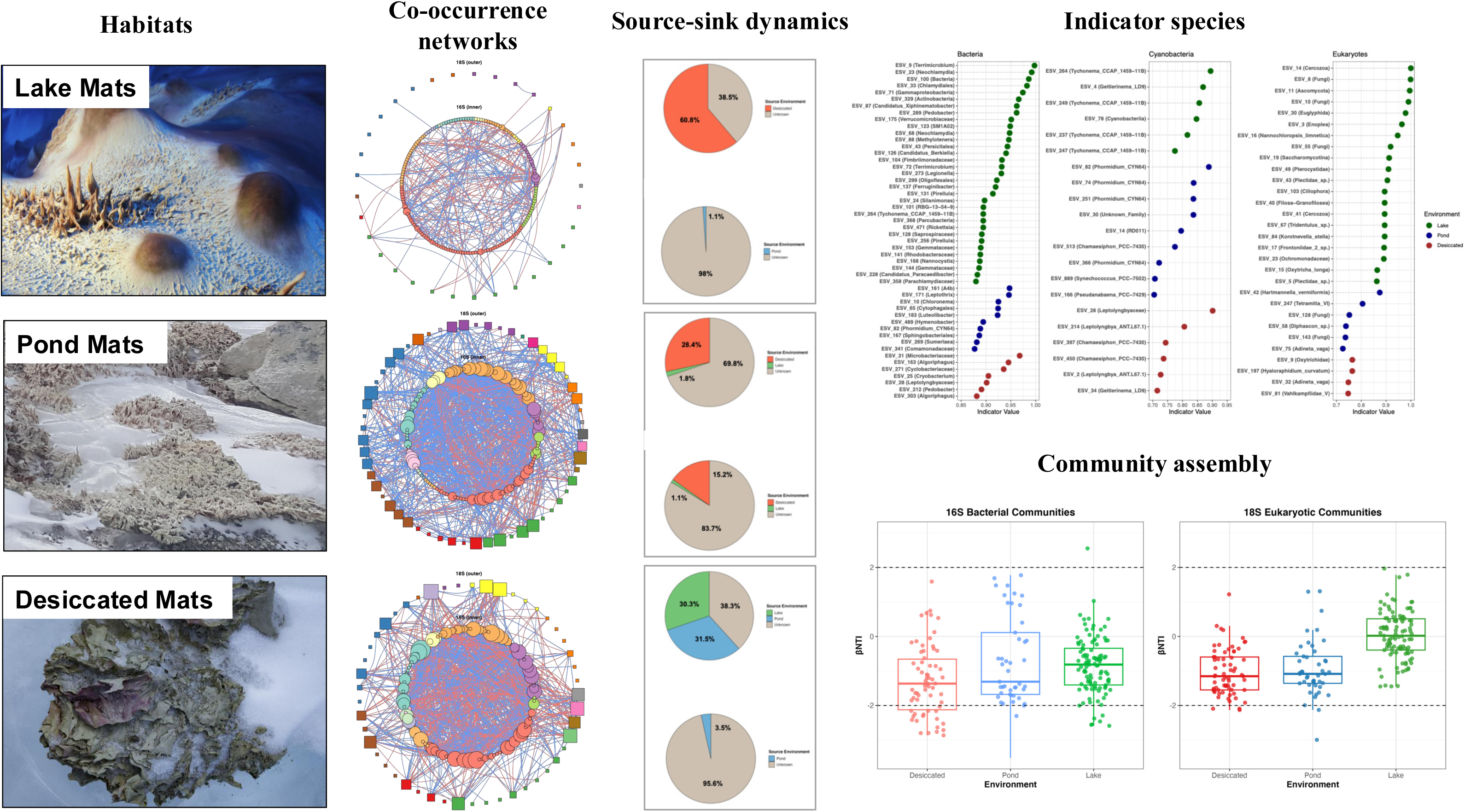

